# Passive surveillance for shrimp pathogens in *Penaeus vannamei* submitted from 3 Regions of Latin America

**DOI:** 10.1101/2023.08.29.555391

**Authors:** Pablo Intriago, Andres Medina, Nicole Cercado, Kelly Arteaga, Alejandra Montenegro, Milena Burgos, Jorge Espinoza, James A Brock, Robins McIntosh, Tim Flegel

## Abstract

Multiple PCR analyzes were performed using 19 different primer sets to open and broaden the search spectrum for shrimp pathogens. In addition, multiple primer pairs for 10 pathogens were compared to see if there were differences in selectivity or sensitivity among them. Some pathogens that did not present histological lesions were detected. The most important outcome was that selection of appropriate primers was the most critical factor in obtaining reliable results. We found high variability in results among primers and we learned it was prudent to seasonally assess among them for the best set selection. It is important to understand that a PCR positive test result alone does not confirm the presence of a viable pathogen or a disease state. Nor, as might be expected, does it mean that the positive PCR test results will be necessarily accompanied by histological lesions characteristic of the targeted pathogen. However, the use of appropriately selected primers sets can reveal whether there is an evolution in the result spectrum over time and if some pathogens disappear or reappear or new ones emerge. In general, most shrimp presented coinfections that consisted of the presence of WzSV8, DHPV, chronic midgut inflammation and tubule distension/epithelial atrophy consistent with Pir A/B toxicity. Also included were RLB/NHPB, microsporidia, striated muscle necrosis, gregarines in the hindgut caecum (gametocyte stage, and not associated with tegumental glands but glands that line the mouth and anus) and encysted, presumed nematode larvae. WzSV8 was newly discovered in gonads. Histological changes and the presence of spheroids in the lymphoid organ were considered as healthy host responses of often unidentified cause.

## 1. INTRODUCTION

### 1.1. General background

Overall shrimp farming production from Latin America has increased yearly during the last decade with the main contribution from Ecuador that in 2022 exported in excess of 1 million MT of shrimp worth over 6.5 billion dollars (**Figure 1**). The exponential increase was due to the adoption of some Asian shrimp culture technology and its very successful adaptation to local conditions. The use of nurseries, aeration, improved pumping capacity and developments in nutrition and feed management (including automatic feeders) have allowed an increase in density and carrying capacity of the systems. The most common stocking density was formerly in the range of 8-12 Postlarvae (PL)/m^2^, while currently, 20 PL/ m^2^ and up to 40 PL/ m^2^ are not unusual.

**Figure 1:**
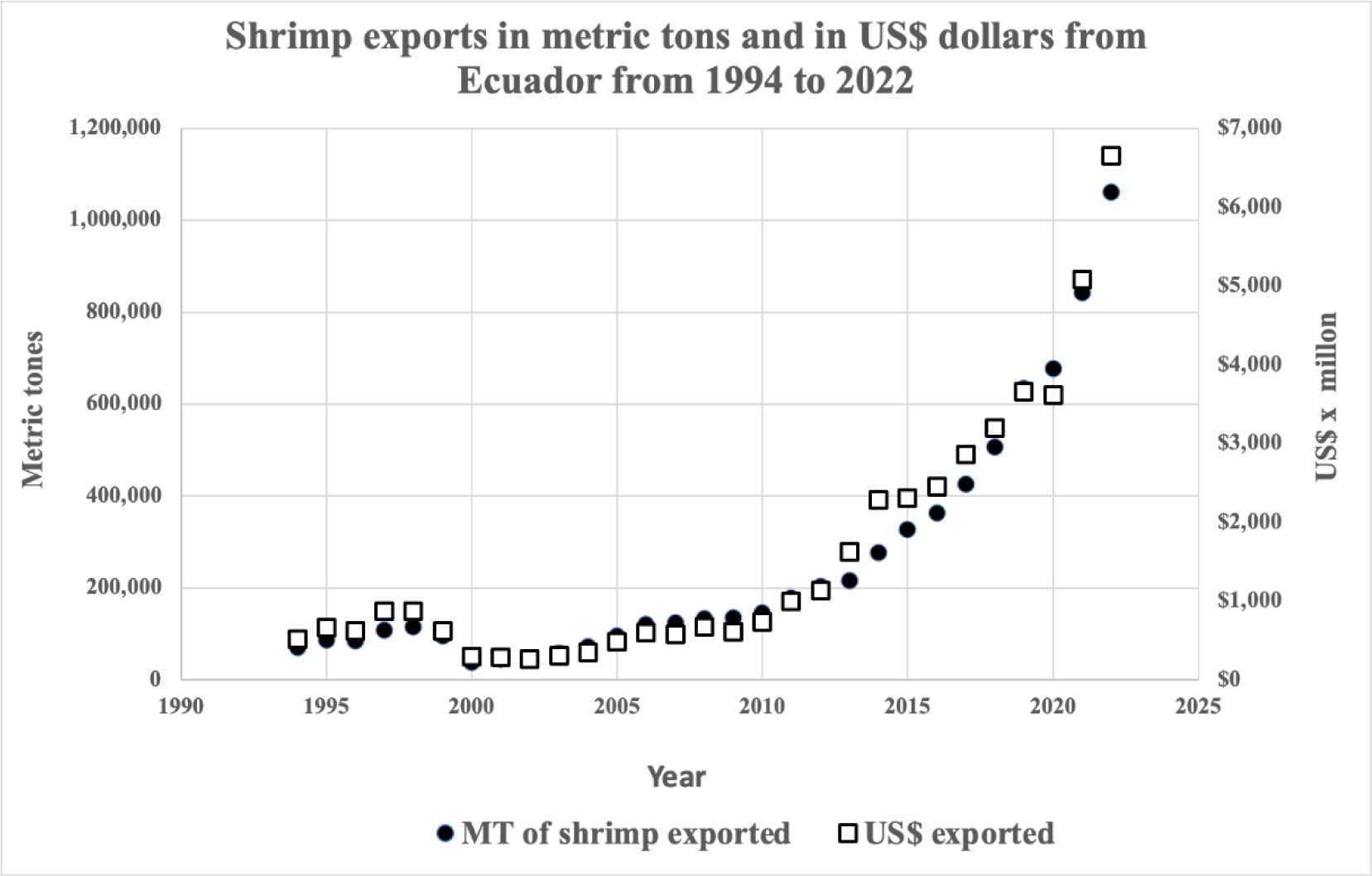
Shrimp exports in metric tons and in US$ dollars from Ecuador from 1994 to 2022.

In Asia specific pathogen free (SPF) broodstock are the dominant source of post larvae (PL) to stock shrimp farms. In contrast, the industry in Latin America relies mostly on pond reared broodstock that present a biosecurity risk for introduction of pathogens into the production cycle. This occurs via pathogen transmission to their offspring PL, perpetuating the presence of pathogens in rearing ponds and potential disease outbreaks. Thus, the current trend for increasing stocking densities and biomass can become excessive and can trigger diseases, especially under unfavorable rearing conditions.

Disease has had a major impact on shrimp aquaculture in the Americas since it became a significant commercial activity in the 1970s (Lightner, 2011). The following pathogens were included in this survey: Infectious hypodermal and haematopoietic necrosis virus (IHHNV)(Lightner et al., 1983; Kalagayan et al. 1991) renamed *Penstylhamaparvovirus* 1 (Pénzes et al, 2020) but referred to herin as IHHNV; Decapod iridescent virus 1 (DIV1)(Qiu et al. 2017); Hepatopancreatic parvovirus (HPV) (Pantoja & Lightner 2000) renamed Decapod hepanhamaparvovirus (DHPV) (Pénzes et al,, 2020) and herein referred to as HPV/DHPV; Macrobrachium hepatopancreatic bidnavirus (MHBV)(Gangnonngiw et al. 2022); Infectious myonecrosis virus (IMNV)) (Lightner et al., 2004a, b); Yellow head virus (YHV) (Tang and Lightner, 1999); Covert mortality nodavirus (CMNV)(Zhang et al. 2014); Penaeus vannamei nodavirus (PvNV)(Poulos & Lightner; 2006); Wenzhou shrimp virus 8 (WzSV8) (Li et al., 2015) later named Penaeus vannamei picornavirus (PvNV) (Liu et al. 2021) followed by Penaeus vannamei solinvivirus (PvSV) (Cruz-Flores et al. 2022) but called WzSV8 herein; Macrobrachium rosenbergii nodavirus (MrNV) (Sri Widada et al., 2003) also known to infect *P. vannamei* (Senapin et al., 2012); the microsporidium *Enterocytozoon hepatopenaei* (EHP) (Tourtip et al., 2009); intracellular bacteria including *Spiroplasma* (Nunan et al. 2004), rickettsia-like bacteria (RLB) (Nunan et al. 2003a,b) and extracellular bacteria including *Vibrio* species (Mohney et al. 1994). All these pathogens have been reported to affect penaeid shrimp culture. In Latin America, some pathogens traditionally considered as virulent to shrimp, such as IHHNV, are rarely found by histological analysis (Jimenez et al. 1999).

The currently most serious shrimp pathogens in Latin America are White spot syndrome virus (WSSV), Taura syndrome virus (TSV), EHP, and the bacteria that cause Acute hepatopancreatic necrosis disease (AHPND) that have collectively cost the penaeid shrimp industry tens of billions of dollars in lost crops, jobs, and export revenue (Lo et al. 1996, Nunan et al., 1998, Shinn et al. 2018, Lightner 2003, Overstreet & Jovonovich 2008, Lightner, 2011, Tran *et al*. 2013).

White spot disease (WSD) caused by WSSV was the dominant disease problem of farmed shrimp in the world until the occurrence of AHPND from 2009-2012 caused by a type of *Vibri*o *parahaemolyticus* carrying a plasmid encoding Pir A/B^VP^ toxins (Lightner *et al*. 2012; Sirikharin et al., 2015). AHPND began to cause significant production losses in Asia (Tran *et al*. 2013) and was subsequently reported from Mexico (Nunan *et al*. 2014; Soto-Rodriguez *et al*. 2015). Since then, other *Vibrio spp.* have been reported to carry this plasmid and cause AHPND (Xiao et al. 2017). The next major pathogen described was hepatopancreatic microsporidiosis (HPM) caused by *Enterocytozoon hepatopenaei* (EHP) first described in 2009 (Tourtip *et al*. 2009) and subsequently reported as a risk factor for APHND in 2014 (Chaijarasphong et al. 2021). Contrary to other pathogens, EHP is not normally associated with mortality but it may impair growth causing severe economic losses depending on the degree of infection.

The use of infected fresh feeds in maturation, the transboundary movement of stocks for farming and the use of pond reared broodstock exposed to wild crustaceans are likely to be the major causes of introducing pathogens into aquaculture systems. Excessive stocking densities and poor management are known factors leading to the expression of diseases (Lightner *et al*. 2012., Tandel *et al*. 2017., Arulmoorthy *et al*. 2020., Tang *et al*. 2020., Albalat *et al*. 2022., Lee *et al*. 2022, Srisala *et al*. 2023). Examples of viruses later discovered in *P. vannamei* include white tail disease in penaeid shrimp caused by infectious myonecrosis virus (IMNV), *Penaeus vannamei* nodavirus (PvNV) (Tang *et al*. 2007b) or by *Macrobrachium rosenbergii* nodavirus (MrNV) (Senapin *et al*. 2012). These 3 viruses target primarily the skeletal muscle and result in very similar gross signs, namely, white, or opaque tail muscles. Histologically, lesions are almost indistinguishable, and in general all of them are characterized by muscle necrosis, hemocytic inflammation and the formation of prominent lymphoid organ spheroids (Tang *et al*. 2007b, Senapin *et al*. 2012). There are no confirmed reports of actual YHV outbreaks in the Americas, YHV has been detected in intensive freshwater cultured *P. vannamei* in Mexico (Sanchez-Barajas et al. 2009). However, there are reports which indicated that YHV infections were co-occurring with white spot disease outbreaks in the USA and in Central America (Pantoja and Lightner. 2003), and an apparently avirulent genotype of YHV is present in farmed and wild penaeid shrimp in northwest Mexico (Lightner, 2011).

Morales-Covarrubias *et al*. (2018) reviewed bacterial diseases in farmed shrimp from 12 regions in Latin America from 2000 to 2015. The most prevalent diseases reported were septic hepatopancreatic necrosis (SHPN), commonly referred to as “vibriosis”, where *V. harveyi, V. parahaemolyticus, V. alginolyticus* and other species were commonly isolated, followed by necrotizing hepatopancreatitis (NHP), streptococcosis, AHPND and spiroplasmosis. The presentation of AHPND in Latin America has been very different to that in Southeast Asia where the major impact was found in ponds (Aranguren Caro *et al*. 2020). In Latin America AHPND has been reported to cause high mortality in *P. vannamei* larviculture and nurseries (Intriago et al, 2023) but not grow out where mortality related to AHPND has been rare and tends to present as a chronic disease (low, sporadic mortality). The difference in the presentation in the ponds might be related to the culture conditions as the origin of AHPND is not an infection but a toxicosis. Environmental conditions (unconsumed feeds, organic load, accumulation of molts, etc.…) determine the replication of the of bacteria and the production of toxin. Latin America practices from low to semi-intensive culture while Asia tends to culture under intensive to super intensive conditions. The difference in the presentation in larviculture and nurseries may also be related to the source of broodstock. While Asia tends to use SPF broodstock as mentioned before, Latin America uses pond reared broodstock that may have brought AHPND back into the production cycle.

More recently, a new shrimp virus was first described in *P. vannamei*. It was initially named Wenzhou shrimp virus 8 (WzSV8) (GenBank record KX883984. 1) from a wide environmental RNA screening of arthropods for viral pathogens (Li *et al*. 2015). In 2018, it was reported in the transcriptome of wild *Penaeus monodon* in Australia (Huerlimann *et al*. 2018). It was subsequently named *Penaeus vannamei* picornavirus (*Pv*PV) when isolated from moribund white leg shrimp (*P. vannamei*) collected a shrimp farm in China in 2015 (Liu et al, 2021). Phylogenetic analysis revealed that *Pv*PV was closely related to WzSV8 (Liu et al., 2021). Srisala et al. (2022, 2023) were the first to identify and characterize WzSV8 lesions in hematoxylin and eosin (H&E) stained tissues examined with a light microscope. Cruz-Flores et al. (2022) reported yet another type of WzSV8 from Brazil with high genome sequence similarity to the sequences of WzSV8 reported from Thailand, China, and Australia. Based on recent changes in viral taxonomy, they named it Penaeus vannamei solinvivirus (PvSV). The impact of PvSV on production of cultivated shrimp has yet to be evaluated.

This paper presents and discusses diagnostic results of over 100 samples processed by histology and PCR. These were collected from 3 different regions in Latin America and from different stages of culture (larvae, broodstock, juvenile/adults) and from the wild. Many of the samples were sent by clients for monitoring shrimp health status or disease outbreaks in ponds. Most of the broodstock were grossly healthy and were collected as reference specimens to assess whether they might be vehicles for introducing pathogens into the production cycle via PL used to stock rearing ponds. Finally, wild shrimp were collected as a reference to what agents might be present as an environmental threat to the industry. More than 20 agents including DNA and RNA viruses, bacteria and microsporidia were targeted. In summary these were:

**DNA viruses**: DHPV, MHBV, DIV1, WSSV and IHHNV.

**RNA viruses**: WzSV8 / PvSV, PvNV, CMNV, IMNV, YHV, TSV and *Mr*NV.

**Bacteria and others**: *Spiroplasma*, *Propionigenium*, Rickettsia Like Bacteria (RLB), Necrotizing Hepatopancreatitis Bacteria (NHP-B), *Vibrio spp*, AHPND, EHP and other non EHP-microsporidia.

Several primers sets were used for some of the microorganisms to test for possible sequence variation in the region and to evaluate the consistency between the PCR and histological results.

### 1.2. Rationale of the study

This study was not an epidemiological study but a simple prevalence analysis of various shrimp pathogens in randomly provided samples from hatcheries, farms, maturation units and wild animals from 3 different regions in Latin America from October 2022 to April 2023. We decided to test different primers because of our experience that histological findings sometimes did not match the PCR results, e.g., the lack of correspondence between histology and PCR for intracellular bacteria, microsporidia and DHPV. We also tested different sets of primers such as those for the spore wall protein (SWP) gene of EHP, because of reports of cross reactions from existing PCR detection methods that target the EHP small subunit ribosomal RNA gene (Jaroenlak et al., 2016; Tang et al., 2017; Dhar et al. 2023). We did tests with DNA from closely related microsporidia and form other aquatic organisms from different regions in Latin America. We also tested different primers of the new virus WzSV8, as there are several reports of it from Asia to America. In this connection, Fredriksson et al. (2013) working with samples from a wastewater plant found that the choice of PCR primers had an impact on assessments of bacterial community and diversity and population dynamics. In addition, Klindworth et al. (2013) found that out of the 175 primers and 512 primer pairs checked, only 10 could be recommended as broad range primers. In conclusion, even commonly used single primers exhibited significant differences in overall coverage and phylum spectrum.

Finally, we were interested in including the detection of *Propionigenium* not as a pathogen or part of the white feces’ syndrome but to test it as biomarker, by presence and copy number, as a measure of the anaerobic or health status of pond bottoms.

## 2. MATERIALS AND METHODS

### 2.1. Sample collection

More than 120 samples from surveillance sampling of *P. vannamei* originating from three different regions in Latin America were analysed throughout the period spanning October 2022 and April 2023. One region draws its culture water from the Pacific Ocean and the other two from the Atlantic Ocean. Sampling when possible included animals from hatcheries, broodstock centres, farms, and wild animals. It should be noted that shrimp sampled for PCR and histology were different individuals from the same populations. To protect client privacy, the countries, or exact locations from which the samples were obtained will not be revealed here. However, the clients from World Organization for Animal Health (WOAH) member countries were informed of their responsibility to notify the competent authority of their country regarding positive test results for any shrimp pathogens listed by WOAH or arising from any unusual incidences of mortality. It would then be the responsibility of the relevant competent authorities from those member countries to report to WOAH.

### 2.2. PCR methods used

DNA was extracted from whole larvae, tissue or organs fixed in 90% alcohol following the manufacturer’s protocol (Omega, Bio-Tek E.Z.N.A tissue DNA kit). In brief, each sample was minced with sterilized scissors and then ground using a microcentrifuge pestle. About 200 mg of tissue was then moved to a clean 1.5 mL Eppendorf tube. To this, 500 μL of tissue lysis buffer (TL) and 25 μL of Omega Biotek (OB) protease solution were added, and the mixture was vortexed and then incubated in a thermoblock at 55°C for approximately 3 h with vortexing every 30 minutes. RNA was removed by adding 4 μL of RNase A (100 mg/mL), and after mixing, the sample was kept at room temperature for 2 min. The sample was then centrifuged at 13,500 RPM for 5 min, and the supernatant was carefully transferred to a new 1.5 mL Eppendorf tube. To this, 220 μL of BL buffer was added, and the mixture was vortexed and incubated at 70°C for 10 mins. Next, 220 μL of 100% ethanol was added, vortexed, and the contents were passed through a HiBind® DNA Mini Column into a 2 mL collection tube. The columns were then centrifuged at 13,500 RPM for 1 min, and the filtrate was discarded. Subsequently, 500 μL of HBC buffer (diluted with 100% isopropanol) was added to the column, and the sample was spun at 13,500 RPM for 30 seconds. The filtrate was discarded, and the column was washed twice with 700 μL of DNA wash buffer diluted with 100% ethanol, and the sample was spun at 13,500 RPM for 30 seconds. The filtrate was discarded. This step was repeated. The column was then centrifuged at 13,500 RPM for 2 mins to dry it out. The dried column was placed in a new nuclease-free 1.5 mL Eppendorf tube, and 100 μL of Elution Buffer, which was heated to 70°C, was added to the column. The sample was allowed to sit for 2 mins before being centrifuged at 13,500 RPM for 1 min. This elution step was repeated. The eluted DNA was then stored at -20°C until needed.

RNA was extracted from whole larvae, tissue or organs fixed in 90% alcohol following the manufacturer’s protocol (Omega, Bio-Tek E.Z.N.A. Total RNA Kit). In brief, each sample was minced with sterilized scissors and then ground using a microcentrifuge pestle. About 200 mg of tissue was then moved to a clean 1.5 mL Eppendorf tube. To this, 700 μL TRK Lysis Buffer was added and the tube was left at room temperature for approximately 3 h with vortexing every 30 minutes. The sample was then centrifuged at 13,500 RPM for 5 mins, and the supernatant was carefully transferred to a new 1.5 mL Eppendorf tube to which 420 μL of 70% ethanol was added. After vortexing to mix thoroughly, the contents were passed through a HiBind® RNA Mini Column into a 2 mL collection tube. The columns were then centrifuged at 13,500 RPM for 1 min, and the filtrate was discarded. Subsequently, 500 μL of RNA Wash Buffer I, was added to the column, and the sample was spun at 13,500 RPM for 30 seconds. The filtrate was discarded, and the column was washed twice with 500 μL RNA Wash Buffer II, diluted with 100% ethanol. The column was then centrifuged at 13,500 RPM for 1 min to dry it out. The filtrate was discarded. This step was repeated. The column was then centrifuged at 13,500 RPM for 2 mins to dry it out. The dried column was placed in a new nuclease-free 1.5 mL Eppendorf tube, and 70 μL of nuclease-free water was added to the column. The sample was centrifuged at 13,500 RPM for 2 min. This elution step was repeated. The eluted RNA was then stored at -70°C until needed. The pathogens tested are listed in **Table 1**. Among the pathogens studied, we also tested the specificity of different primers (see **Table 2**).

**Table 1.**
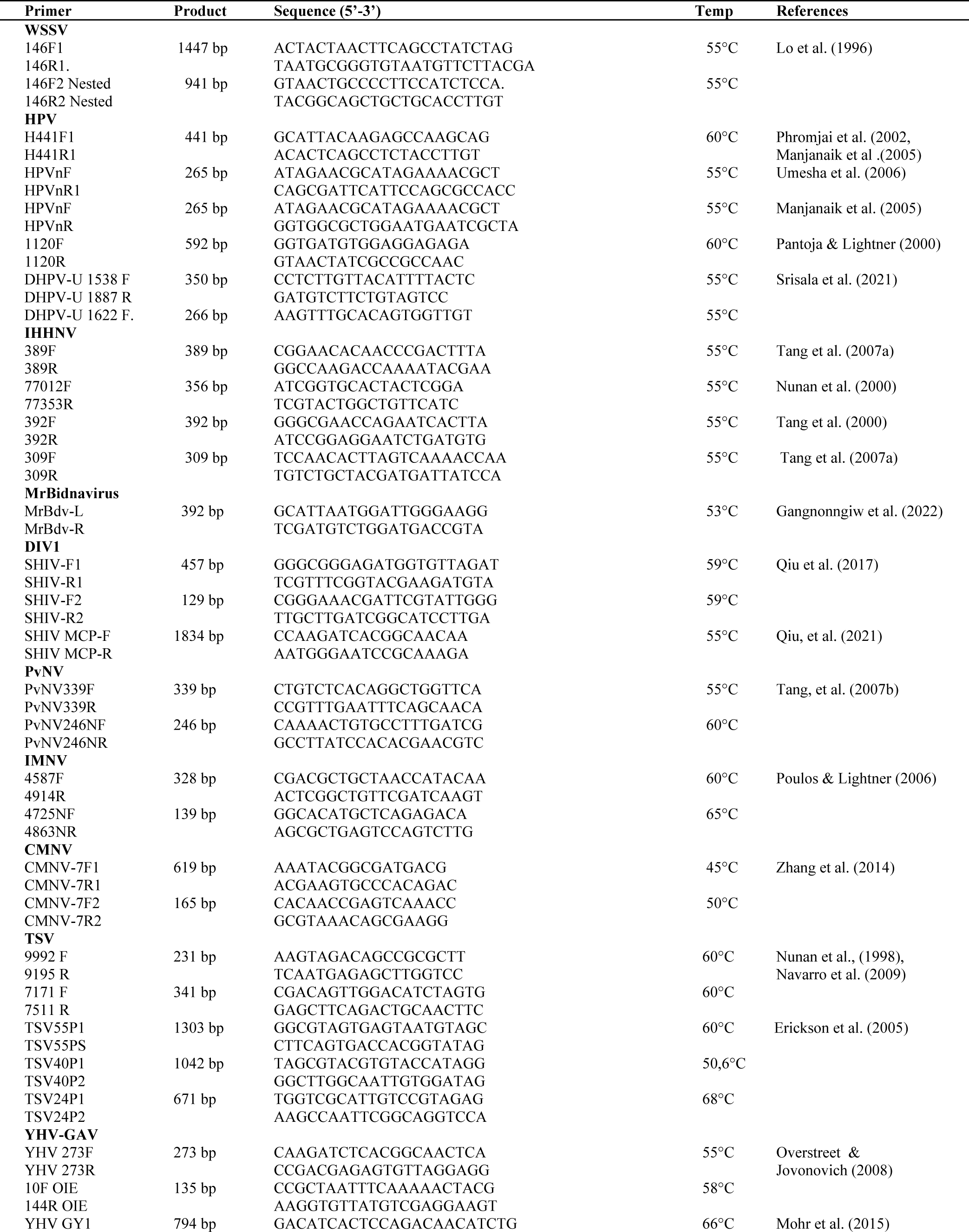
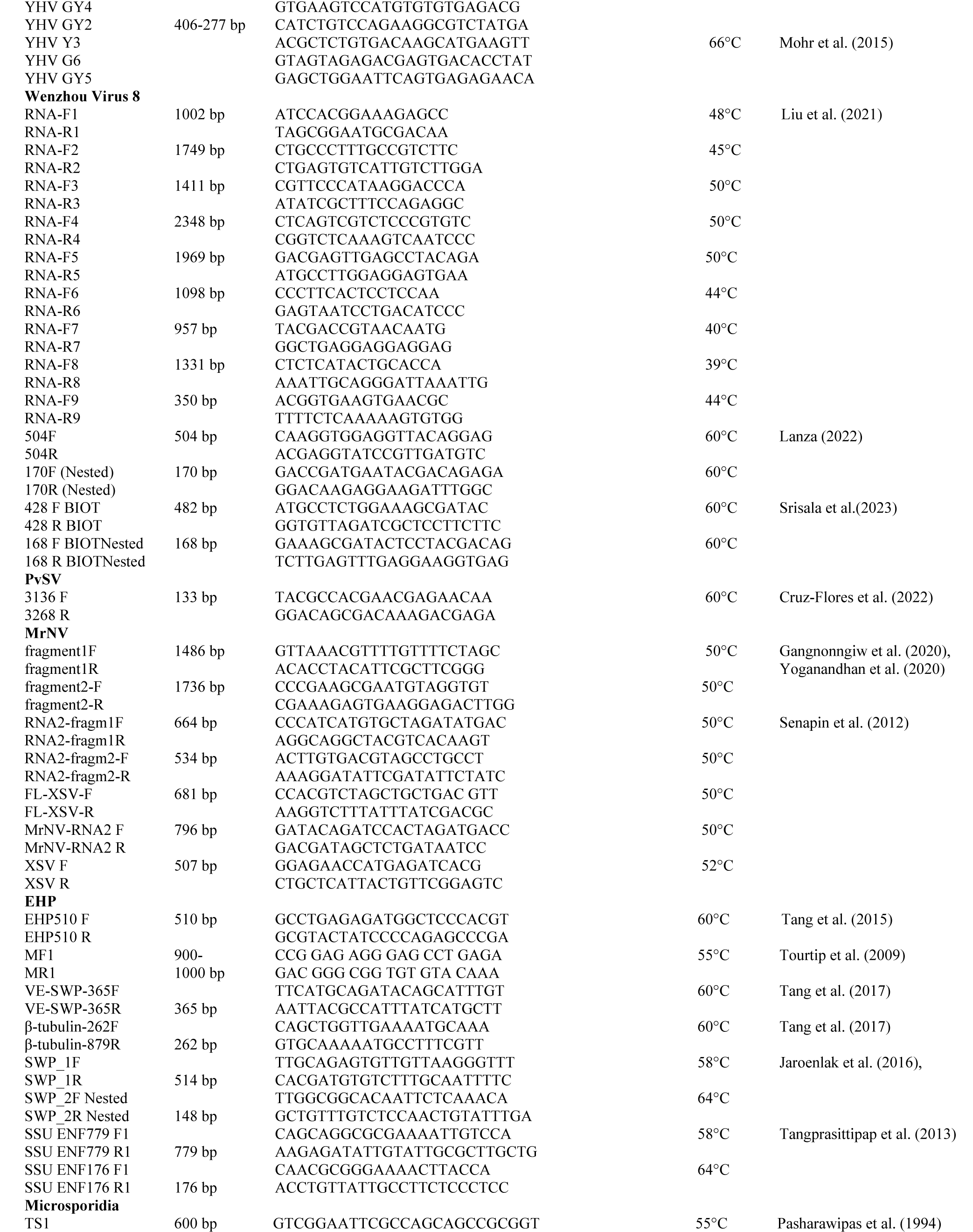
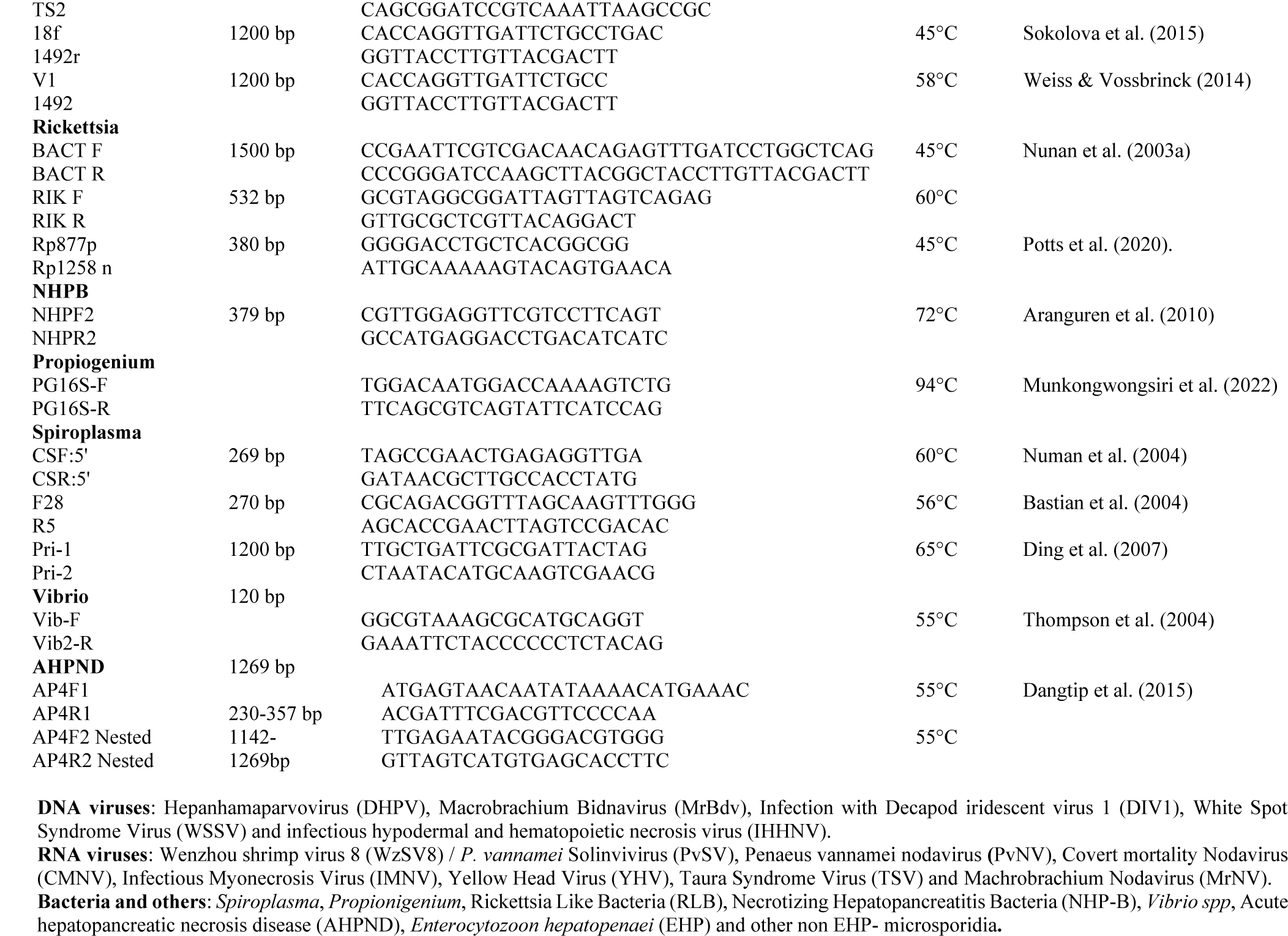
Primer sets used in this study.

**Table 2.** Set of different primers used to test specificity in some pathogens.

| Pathogen |  |  |  |  |  |  |
| --- | --- | --- | --- | --- | --- | --- |
| **HPV- MrBdv-L/R** | H441F/R<br>HPVnF/R nested<br>Phromjai et al.<br>(2002).<br>Umesha et al. (2006) | H441F/R<br>HPVnF/R nested<br>(Phromjai et al.<br>(2002).<br>Srisala et al.<br>(2021) | 1120F/R<br>(Pantoja & Lightner,<br>2000) | DHPV-U 1538<br>F/ 1887 R/ 1622<br>F Semi-Nested<br>Srisala, J.<br>(2021)<br>Manjanaik et al.<br>(2005) | MrBdv-L/R<br>Bidnavirus (MrBdv)<br>Gangnonngiw et al.<br>(2022) |  |
| **DIV1** | Nested SHIV-F/R2<br>Qiu, et al. (2017) | 1834bp<br>Qiu, et al. (2021) |  |  |  |  |
| **IHHNV** | 309 F/R<br>Tang and Lightner,<br>(2006).<br>Tang et al. (2007a) | 392 F/R<br>Tang et al. (2000) | 389 F/R<br>Tang and Lightner,<br>(2006).<br>Tang et al. (2007a) | 77012F/77353R<br>(356bp)<br>Nunan et al.<br>(2000) |  |  |
| **WZ8V/**<br>**PvSV** | WSV8<br>504F/R<br>170F/R<br>Brazil (Lanza, 2022*) | WSV8<br>428F/R-168F/R<br>(Nested)<br>(Srisala et al.<br>2023) | Solinvivirus (PvSV)<br>Cruz-Flores, et al.<br>(2022) | Liu et al. (2021) |  |  |
| **TSV** | TSV (Nunan, et al. 1998); Navarro, et al. 2009)<br>9992/9195-7171/7511 |  | TSV (Erickson et al., 2005) VP1-VP2-VP3 |  |  |  |
| **MrNV** | RNA1-fragment1-F/R<br>RNA1-fragment2-F/R<br>RNA2-fragment1-F/R<br>RNA2-fragment2-F/R |  |  | MrNV2F/R<br>FL-XSV-F/R<br>XSV-F/R |  |  |
| **Spiroplasma**<br>**spp.** | CSF/R<br>Nunan et al. (2004) | F28/R5<br>Bastian et al.<br>(2004) | Pri-1/2<br>Ding et al. (2007) |  |  |  |
| **Intracellular**<br>**bacteria** | Bact F/R<br>Rik F/R<br>Nunan, et al.<br>(2003a) | Rp877p/Rp1258n<br>Potts et al. (2020) | NHPB-FW/RV<br>Aranguren et al.<br>(2010) |  |  |  |
| **EHP** | EHP 510 F/R<br>bp<br>Tang et al. (2015) | EHP Genome 900-1000bp<br>Tourtip et al.<br>(2009) | VE-EHP-SWP-365bp<br>Tang et al. (2017) | SSU<br>Tangprasitti<br>pap et al.<br>(2013) | SWP 1F/1R.<br>SWP 2F/2R<br>Jaroenlak et al.<br>(2016) | EHP B-tubulin<br>Tang et al.<br>(2017) |
| **Microsporidia** | TS1/TS2<br>Pasharawipas et al.<br>(1994) | 18f/ 1492r<br>Sokolova et al.<br>(2015) | Ss2 18f3/ ss1492r<br>Weiss & Vossbrinck (2014) |  |  |  |

Note:
DNA virus: HPV or DHPV, Bidnavirus (MrBdv), Infection with Decapod iridescent virus 1 (DIV1), IHHNV.
RNA virus: Wenzhou shrimp virus 8 (WZSV8) / *P. vannamei* Solinvivirus (PvSV), (Liu et al. 2021, Srisala et al. 2023; Cruz Flores et al. 2022), Brazil sequence was provided by Lanza, 2022, from Universidade Federal do Rio Grande do Norte, Brazil).
Taura Syndrome Virus (TSV) and *Macrobrachium* Nodavirus (MrNV).
Others: *Spiroplasma Penaei* (Nunan et al. 2004), *S mirium* (Bastian et al. 2004), Universal primers for *Spiroplasma* (Ding et al. 2007).
\*Personal communication.

### 2.3. Samples used for extraction

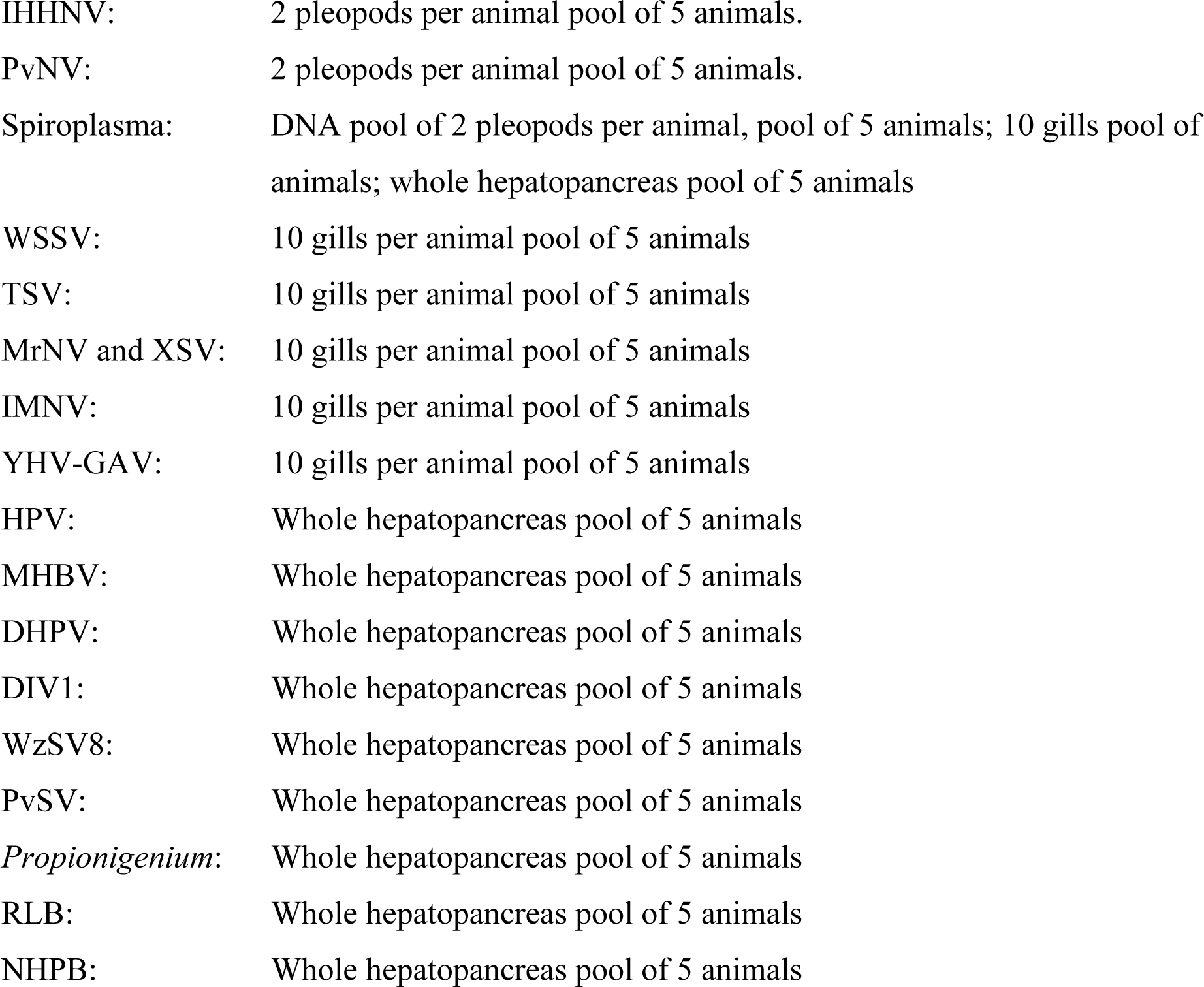
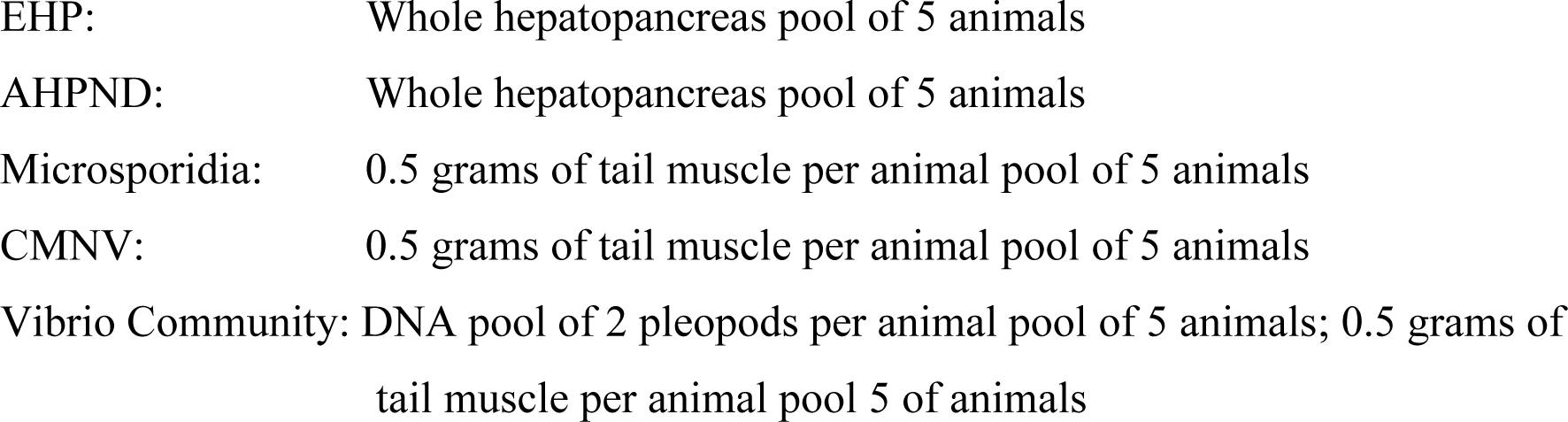

**2.3. Samples used for extraction.**
|  |  |
| --- | --- |
| IHHNV: | 2 pleopods per animal pool of 5 animals. |
| PvNV: | 2 pleopods per animal pool of 5 animals. |
| Spiroplasma: | DNA pool of 2 pleopods per animal, pool of 5 animals; 10 gills pool of animals; whole hepatopancreas pool of 5 animals |
| WSSV: | 10 gills per animal pool of 5 animals |
| TSV: | 10 gills per animal pool of 5 animals |
| MrNV and XSV: | 10 gills per animal pool of 5 animals |
| IMNV: | 10 gills per animal pool of 5 animals |
| YHV-GAV: | 10 gills per animal pool of 5 animals |
| HPV: | Whole hepatopancreas pool of 5 animals |
| MHBV: | Whole hepatopancreas pool of 5 animals |
| DHPV: | Whole hepatopancreas pool of 5 animals |
| DIV1: | Whole hepatopancreas pool of 5 animals |
| WzSV8: | Whole hepatopancreas pool of 5 animals |
| PvSV: | Whole hepatopancreas pool of 5 animals |
| *Propionigenium*: | Whole hepatopancreas pool of 5 animals |
| RLB: | Whole hepatopancreas pool of 5 animals |
| NHPB: | Whole hepatopancreas pool of 5 animals |
| EHP: | Whole hepatopancreas pool of 5 animals |
| AHPND: | Whole hepatopancreas pool of 5 animals |
| Microsporidia: | 0.5 grams of tail muscle per animal pool of 5 animals |
| CMNV: | 0.5 grams of tail muscle per animal pool of 5 animals |
| Vibrio Community: | DNA pool of 2 pleopods per animal pool of 5 animals; 0.5 grams of tail muscle per animal pool 5 of animals |

### 2.4. Histopathology

For histological analysis, samples were prepared following the procedures outlined by Bell and Lightner (1988). Briefly, they were fixed in Davidson’s AFA for at least 24 h or 72 h in case of broodstock before processing for routine histological analysis of 5 µm thick tissue sections stained with hematoxylin and eosin (H&E). From each juvenile to broodstock shrimp sample, 2 to 4 paraffin blocks were prepared. For post-larvae (PL), approximately 1,500 animals were taken from each tank and more than half (i.e., about 750+) were fixed in Davidson’s AFA for 24 h and embedded in paraffin blocks, each containing approximately 35 to 50 PL.

## 3. RESULTS

### 3.1. Specificity and sensitivity of the different PCR primers and protocols

PCR results are summarized in the tables below comparing the specificity and sensitivity of different primers used for WzSV8, DHPV, RLB/NHPB, *Spiroplasma,* Microsporidia, IHHNV, EHP, YHV, TSV, MrNV/XSV, DIV1 for the 3 regions sampled.

Both WzSV8 primers 170 F/R and PvSV (Lanza, 2022 personal communication and Cruz Flores et al. 2022) were the more sensitive (77% and 84% respectively) than the primer designed by BIOTEC (Thailand) (Srisala et al. 2023) with 64% prevalence (**Table 3**). These primers have been developed by American research teams using local isolates. We also unsuccessfully tested all 9 primers listed by Liu and co-authors (Liu et al. 2021) in 28 samples from region 1, after repeated negative results we stopped using these primers.

**Table 3.** Differences found when using 3 different primers to detect presence of WzSV8/PvSV in *P. vannamei* in samples from 3 different regions of Latin America.

| Region/ Primer | Pathogen/s | WzSV8<br>504F/R<br>170F/R<br>Brazil (Nested)* | WzSV8<br>428F/R<br>168F/R<br>BIOTEC<br>(Nested)** | Solinvivirus (PvSV)***<br>Cruz-Flores, et al.<br>(2022) |
| --- | --- | --- | --- | --- |
| Region #1 | WzSV8V |  |  |  |
| PL |  | 6/6 (100%) | 3/6 (50%) | 6/6 (100%) |
| Farm |  | 34/40 (85%) | 25/40 (63%) | 36/36 (100%) |
| Broodstock |  | 17/24 (71%) | 16/24 (67%) | 18/24 (75%) |
| Wild Broodstock |  | 2/12 (17%) | 5/12 (42%) | 9/12 (75%) |
| Region #2 |  |  |  |  |
| PL |  | 0/1 (0%) | 0/1 (0%) | 1/1 (100%) |
| Farm |  | 17/17 (100%) | 11/17 (65%) | 12/17 (71%) |
| Region #3 |  |  |  |  |
| Farm |  | 15/18 (83%) | 15/18 (83%) | 14/18 (78%) |
| Average <sup>1</sup> |  | 91/118 (77%) | 75/118 (64%) | 96/114 (84%) |

Lanza (2022) personal communication \*. Srisala et al. (2023) \*\*. Cruz-Flores, et al. (2022) \*\*\*.<sup>4</sup> Liu et al (2021).
<sup>1</sup> Average pool of all samples per primer in all regions.

**Table 4** shows a comparison of 4 different primers used to detect DHPV plus MHBV. The nested method used by Phromjai et al. (2002) and Umesha et al. (2006) and a later modified version (Phromjai et al. 2002, Srisala et al. 2021), were more sensitive (14 and 12% respectively) when compared to other methods including the semi nested method reported as universal DHPV (5% prevalence) (Srisala et al. 2022) and the 1120F/R from Pantoja & Lightner (2000) with 3% prevalence. MHBV was always negative and was included because of a previous report that its intranuclear inclusion bodies are somewhat like those of DHPV types (Gangnonngiw et al. 2022), while the type of DHPV reported from *M. rosenbergii* does not produce intranuclear inclusions.

**Table 4.**
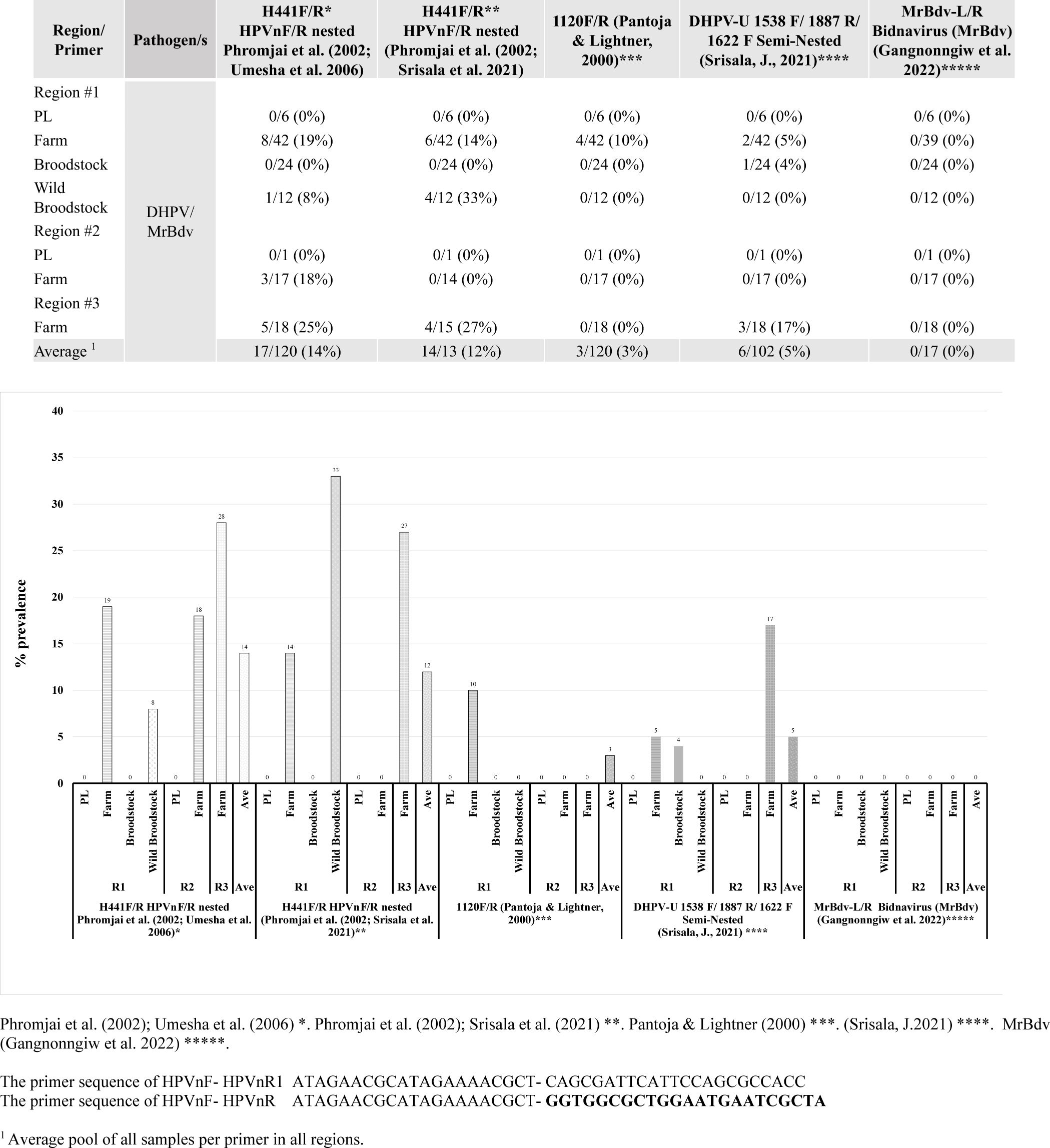
Differences found when using 4 different primers to detect presence of DHPV in *P. vannamei*. Primers to detect MrBdv were included as comparison between these two viral pathogens. Samples were from 3 different regions of Latin America.

Interestingly, the detection of WzSV8 and DHPV in wild broodstock in region 1 showed different patterns. On one hand, WzSV8 170 F/R was less sensitive (17% prevalence) than 428F/R-168F/R from Srisala et al. (2023) (42% prevalence) and PvSV (Cruz-Flores et al. 2022) (75% prevalence). On the other, the modified version of DHPV of Phromjai et al. 2002, Srisala et al. (2021) was more sensitive (33% vs 8%) than the primers adapted by Phromjai et al. (2002) and Umesha et al. (2006).

In the present study Rickettsia like bacteria were the most common type of intracellular bacteria detected, as RLB (Nunan et al. 2003a, b) or detected with a universal primer (Potts et al. 2020) (28% and 25% respectively), with a significant prevalence in all three regions. The prevalence of NHPB was very low (1% overall) and was found in only one sample in region 3 (**Table 5**).

**Table 5:**
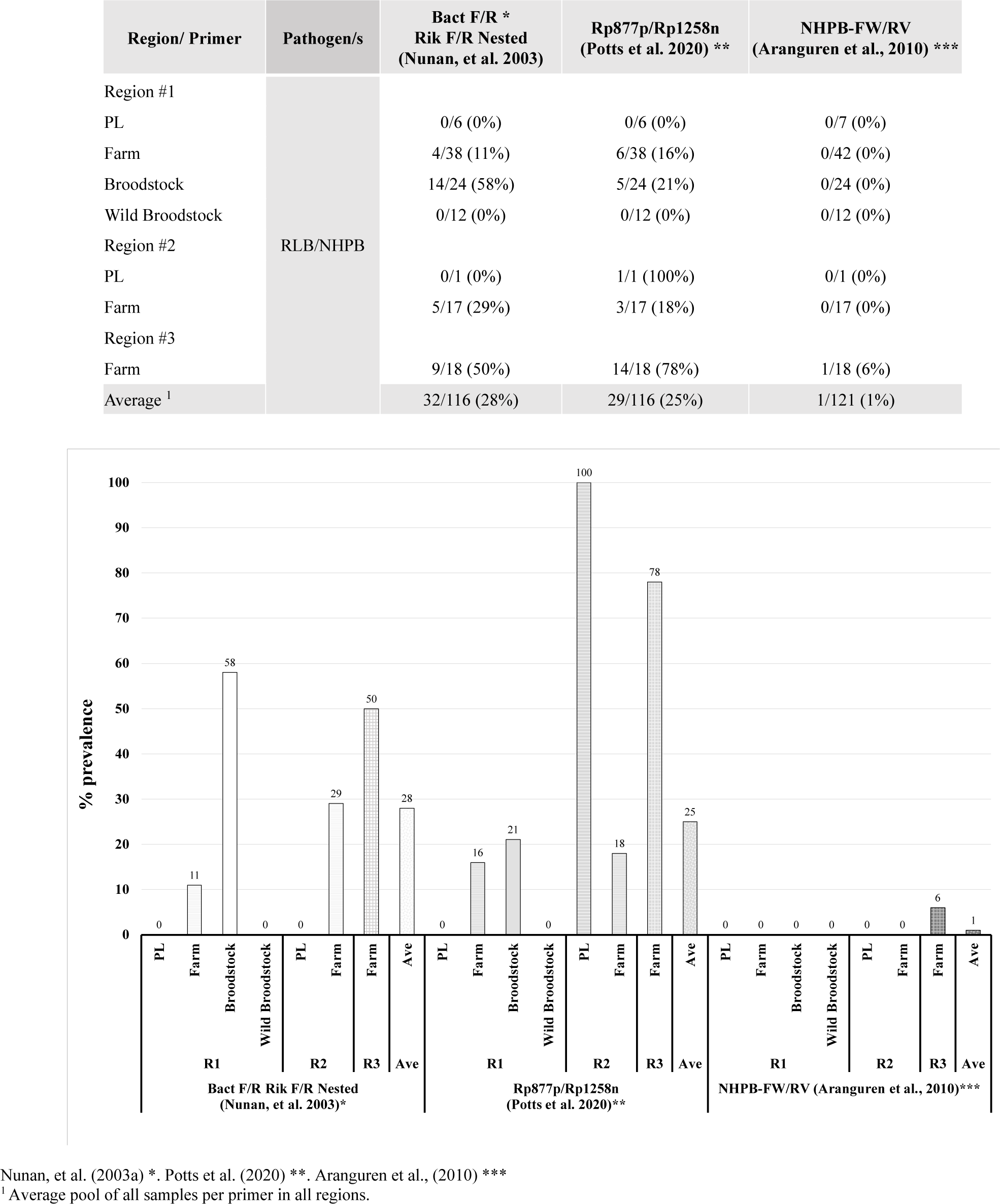
Differences found when using different primers to detect presence of intracellular bacteria in *P. vannamei* in samples from 3 different regions of Latin America.

| Region/ Primer | Pathogen/s | Bact F/R *<br>Rik F/R Nested<br>(Nunan, et al. 2003) | Rp877p/Rp1258n<br>(Potts et al. 2020) ** | NHPB-FW/RV<br>(Aranguren et al., 2010) *** |
| --- | --- | --- | --- | --- |
| Region #1 | RLB/NHPB |  |  |  |
| PL |  | 0/6 (0%) | 0/6 (0%) | 0/7 (0%) |
| Farm |  | 4/38 (11%) | 6/38 (16%) | 0/42 (0%) |
| Broodstock |  | 14/24 (58%) | 5/24 (21%) | 0/24 (0%) |
| Wild Broodstock |  | 0/12 (0%) | 0/12 (0%) | 0/12 (0%) |
| Region #2 |  |  |  |  |
| PL |  | 0/1 (0%) | 1/1 (100%) | 0/1 (0%) |
| Farm |  | 5/17 (29%) | 3/17 (18%) | 0/17 (0%) |
| Region #3 |  |  |  |  |
| Farm |  | 9/18 (50%) | 14/18 (78%) | 1/18 (6%) |
| Average <sup>1</sup> |  | 32/116 (28%) | 29/116 (25%) | 1/121 (1%) |

Nunan, et al. (2003a) \*. Potts et al. (2020) \*\*. Aranguren et al., (2010) \*\*\*
<sup>1</sup> Average pool of all samples per primer in all regions.

Nunan et al. (2004) were the first to report *Spiroplasma* as a pathogen to *P. vannamei* shrimp from samples with severe mortalities at a Colombian shrimp farm. Nunan et al. (2004) named this species *S. penaei* and designed a set of primers for its detection. **Table 6** shows the prevalence of this pathogen as well as two other different primers reported for *Spiroplasma*-specific 16S rDNA, namely F28/R5 (Bastian et al. 2004), and Pri 1/2 (Ding et al. 2007). Both primers have been reported to detect *S. mirium* as well as *S. eriocheiris* (Ding et al. 2007, Liang et al. 2010). Interestingly, besides *S. penaei* (24% prevalence), Pri 1/2 primers also detected (10% prevalence) in regions 1 and 3. Ding et al. (2007) developed this set for detecting *Spiroplasma* in the Chinese mitten crab *Eriocheir sinensis*, and the crayfish *Procambarus clarkii*. They found that both Pri 1/2 and F28/R5 detected it in some samples, a result that differs from those from the present study.

**Table 6:** Differences found when using 3 different primers to detect presence of *Spiroplasma* spp. in *P. vannamei* in samples from 3 different regions of Latin America.

| Region/ Primer | Pathogen/s | CSF/R *<br>(Nunan et al. 2004) | F28/R5 **<br>(Bastian et al. 2004) | Pri-1/2 ***<br>(Ding et al. 2007) |
| --- | --- | --- | --- | --- |
| Region #1 | Spiroplasma |  |  |  |
| PL |  | 2/7 (29%) | 0/7 (0%) | 0/6 (0%) |
| Farm |  | 14/37 (38%) | 0/37 (0%) | 3/37 (8%) |
| Broodstock |  | 0/24 (0%) | 0/24 (0%) | 0/24 (0%) |
| Wild Broodstock |  | 0/12 (0%) | 0/12 (0%) | 0/12 (0%) |
| Region #2 |  |  |  |  |
| PL |  | 1/1 (100%) | 0/1 (0%) | 0/1 (0%) |
| Farm |  | 0/17 (0%) | 0/17 (0%) | 0/17 (0%) |
| Region #3 |  |  |  |  |
| Farm |  | 7/18 (39%) | 0/18 (0%) | 8/18 (44%) |
| Average <sup>1</sup> |  | 24/116 (21%) | 0/116 (0%) | 11/115 (10%) |

Nunan, et al. (2004) \*. Bastian et al. (2004) \*\*. Ding et al. (2007) \*\*\*
<sup>1</sup> Average pool of all samples per primer in all regions.

For microsporidia in the present study, we tested two set of universal primers 18f and 1492r used by Sokolova et al. (2015) and (Vossbrinck. et al., 2004), which amplify most of the small subunit rRNA of microsporidia. We included Pasharawipas et al. (1994) primers that are selective for *A. penaei* (**Table 7**). *A. penaei* primers were more sensitive than both universals tested (50% vs 36% and 19%). Microsporidia were detected in farms from three regions. Farm samples of region 3 had the highest prevalence with 94% of the samples positive.

**Table 7.** Comparison of 3 different primers to detect presence of Microsporidia in *P. vannamei* in samples from 3 different regions of Latin America.

| Region/ Primer | Pathogen/s | TS1/TS2<br>(Pasharawipas et al. 1994) * | 18f/ 1492r<br>(Sokolova et al. 2015) ** | Ss2 18f3/ ss1492r<br>Universal (Weiss & Vossbrinck,<br>2014) *** |
| --- | --- | --- | --- | --- |
| Region #1 | Microsporidia |  |  |  |
| PL |  | 2/6 (33%) | 0/6 (0%) | 0/6 (0%) |
| Farm |  | 22/40 (55%) | 7/40 (18%) | 15/32 (47%) |
| Broodstock |  | 8/24 (33%) | 0/24 (0%) | 2/24 (8%) |
| Wild Broodstock |  | 2/12 (17%) | 5/12 (42%) | 9/12 (75%) |
| Region #2 |  |  |  |  |
| PL |  | 0/1 (0%) | 0/1 (0%) | ND* |
| Farm |  | 8/17 (47%) | 6/17 (35%) | 3/14 (21%) |
| Region #3 |  |  |  |  |
| Farm |  | 17/18 (94%) | 5/18 (28%) | 11/18 (61%) |
| Average <sup>1</sup> |  | 59/118 (50%) | 23/118 (19%) | 40/110 (36%) |

Pasharawipas et al. (1994) \*. Sokolova et al. (2015) \*\*. Weiss & Vossbrinck, (2014) \*\*\*
<sup>1</sup> Average pool of all samples per primer in all regions.

**Table 8** shows a comparison of 4 primers commonly used for IHHNV (309 F/R, 389 F/R. 392 F/R and 77012F/77353R). IHHNV was only found in farm samples of region 1, and within these samples only 15% could be considered as IHHNV complete virus, whereas 23% of the samples were EVEs positive. Both 392F/R and 389F/R were the most common primers followed by 309 F/R and 77012F/77353R.

**Table 8.** Comparison and interpretation when using 4 different set of primers to detect presence of IHHNV in *P. vannamei* in samples from 3 different regions of Latin America.

| Region/ Primer | Pathogen/s | 309 F/R *<br>(Tang and Lightner,<br>20006, Tang et al. 2007a) | 392 F/R **<br>(Tang et al.<br>2000) | 389 F/R ***<br>(Tang and Lightner,<br>2006; Tang et al. 2007a) | 77012F/77353R (356bp)<br>(Nunan et al., 2000)<br>**** | IHHNV | eve |
| --- | --- | --- | --- | --- | --- | --- | --- |
| Region #1 | IHHNV |  |  |  |  |  |  |
| PL |  | 0/6 (0%) | 0/6 (0%) | 0/6 (0%) | 0/6 (0%) | 0/6 (0%) | 0/6 (0%) |
| Farm |  | 9/40 (23%) | 14/40 (35%) | 12/40 (30%) | 9/40 (23%) | 6/40 (15%) | 9/40 (23%) |
| Broodstock |  | 0/24 (0%) | 0/24 (0%) | 0/24 (0%) | 0/24 (0%) | 0/24 (0%) | 0/24 (0%) |
| Wild Broodstock |  | 0/12 (0%) | 0/12 (0%) | 0/12 (0%) | 0/12 (0%) | 0/12 (0%) | 0/12 (0%) |
| Region #2 |  |  |  |  |  |  |  |
| PL |  | 0/1 (0%) | 0/1 (0%) | 0/1 (0%) | 0/1 (0%) | 0/1 (0%) | 0/1 (0%) |
| Farm |  | 0/17 (0%) | 0/17 (0%) | 0/17 (0%) | 0/17 (0%) | 0/17 (0%) | 0/17 (0%) |
| Region #3 |  |  |  |  |  |  |  |
| Farm |  | 0/18 (0%) | 0/18 (0%) | 0/18 (0%) | 0/18 (0%) | 0/18 (0%) | 0/18 (0%) |
| Average <sup>1</sup> |  | 9/118 (8%) | 14/118 (12%) | 12/118 (10%) | 9/118 (8%) | 6/118 (5%) | 9/118 (8%) |

EVE: When tested using multiple PCR primers, they all must be present for a functional virus to replicate. Whereas, if some but not all primers amplify, this pattern is a clear indication that the PCR primers are measuring an eve (endogenous viral element) and not the complete virus.
Tang and Lightner (2006) \*, Tang et al. (2007a) \*\*. Tang et al. (2000) \*\*\*. Tang and Lightner (2006), Tang et al. (2007a) \*\*\*\*. Nunan et al., (2000) \*\*\*\*\*.
<sup>1</sup> Average pool of all samples per primer in all regions.

**Table 9** shows the results from 6 different primers used for detecting EHP. In general, most of the existing PCR detection methods target the EHP small subunit ribosomal RNA (SSU rRNA) gene (SSU-PCR) (Tourtip et al. 2009, Tangprasittipap et al. 2013, Tang et al. 2015, Tang et al. 2017). However, Jaroenlak et al. (2016) discovered that they can give false positive test results due to cross reactivity of the SSU-PCR primers with DNA from closely related microsporidia. To overcome this problem, a nested PCR method was developed for detection of the spore wall protein (SWP) gene of EHP (Jaroenlak et al. 2016). Only one of our samples from a farm in region 1 gave positive results using the primers targeting the SWP gene, equivalent to an average of 2% of prevalence in this region and 1% overall. The other nested method (Tangprasittipap et al. 2013) gave 2% of prevalence, both in only one sample each from farms in regions 1 and 3. The method of Tang (Tang et al. 2015) gave 3% for overall prevalence from one sample each from different farm samples in the 3 regions. We did not find any positive reactors using the primers of Tang et al (2017). The most abundant prevalence was found using Tourtip et al. (2009) primers with 10% overall prevalence (one sample positive in each of regions 1 and 2 but 10 in 18 samples positive from region 3).

**Table 9.** Comparison between 6 different set of primers to detect presence of EHP in *P. vannamei* in samples from 3 different regions of Latin America.

| Region/ Primer | Pathogen/s | EHP 510bp * | EHP 900-1000bp ** | VE-EHP-SWP-365bp *** | EHP B-tubulin 262 bp *** | SSU ENF779 F1<br>SSU ENF779 R1<br>SSU ENF176 F1<br>SSU ENF176 R1<br>779 bp 176 bp **** | SWP_1F SWP_1R<br>SWP_2F SWP_2R<br>514 bp 148 bp ***** |
| --- | --- | --- | --- | --- | --- | --- | --- |
| Region #1 | EHP |  |  |  |  |  |  |
| PL |  | 0/6 (0%) | 0/6 (0%) | 0/6 (0%) | 0/6 (0%) | 0/6 (0%) | 0/6 (0%) |
| Farm |  | 1/42 (2%) | 0/42 (0%) | 0/42 (0%) | 0/42 (0%) | 1/42 (2%) | 1/42 (2%) |
| Broodstock |  | 0/24 (0%) | 1/24 (4%) | 0/24 (0%) | 0/24 (0%) | 0/24 (0%) | 0/24 (0%) |
| Wild Broodstock |  | 0/12 (0%) | 0/12 (0%) | 0/12 (0%) | 0/12 (0%) | 0/12 (0%) | 0/12 (0%) |
| Region #2 |  |  |  |  |  |  |  |
| PL |  | 0/1 (0%) | 0/1 (0%) | 0/1 (0%) | 0/1 (0%) | 0/1 (0%) | 0/1 (0%) |
| Farm |  | 1/17 (6%) | 1/17 (0%) | 0/17 (0%) | 0/17 (0%) | 0/17 (0%) | 0/17 (0%) |
| Region #3 |  |  |  |  |  |  |  |
| Farm |  | 1/18 (6%) | 10/18 (56%) | 0/18 (0%) | 0/18 (0%) | 1/18 (6%) | 0/18 (0%) |
| Average <sup>1</sup> |  | 3/120 (3%) | 12/120 (10%) | 0/120 (0%) | 0/120 (0%) | 2/120 (2%) | 1/120 (1%) |

Tang et al (2015) \*. Tourtip et al. (2009) \*\*. Tang et al. (2017) \*\*\*. Tangprasittipap et al. (2013) \*\*\*\*. Jaroenlak et al. (2016) \*\*\*\*\*.
<sup>1</sup> Average pool of all samples per primer in all regions.

Results for DIV1, YHV, TSV, YHV-GAV, PvNV, IMNV, MrNV and XSV were not included because they were very rare or absent.

### 3.2. PCR results by region

In general, WzSV8 was the dominant agent in all regions (89%) and was the only one found in all samples independent of region or sizes (**Table 10**). It was followed by *Propionigenium* and microsporidia (75% and 56%) respectively. However, *Propionigenium* is not considered a pathogen but was used in this study as a bio-indicator (**Table 11**). The prevalence of this anaerobic bacterium was 100% in both broodstock and wild animals from region 1. The only group of animals that had no positive test reaction were post larvae from region 2. The highest prevalence of microsporidia was in farm animals (94%) of region 3.

**Table 10:** Prevalence of several pathogens tested for region.

| Region / Pathogen | Spiroplasma | Propionigenium | RLB | Vibrio | AHPND | Microsporidia |
| --- | --- | --- | --- | --- | --- | --- |
| Region #1 |  |  |  |  |  |  |
| PL | 2/7 (29%) | 4/6 (67%) | 0/6 (0%) | 0/6 (0%) | 0/6 (0%) | 2/6 (33%) |
| Farm | 14/37 (38%) | 27/42 (64%) | 6/38 (16%) | 9/38 (24%) | 9/42 (21%) | 22/40 (55%) |
| Broodstock | 0/24 (0%) | 24/24 (100%) | 14/24 (58%) | 9/24 (38%) | 1/24 (4%) | 8/24 (33%) |
| Wild Broodstock | 0/12 (0%) | 12/12 (100%) | 0/12 (0%) | 0/12 (0%) | 0/12 (0%) | 9/12 (75%) |
| Region #2 |  |  |  |  |  |  |
| PL | 1/1 (100%) | 0/1 (0%) | 1/1 (100%) | 0/1 (0%) | 0/1 (0%) | 0/1 (0%) |
| Farm | 0/17 (0%) | 9/17 (53%) | 5/17 (29%) | 2/17 (12%) | 0/17 (0%) | 8/17 (47%) |
| Region #3 |  |  |  |  |  |  |
| Farm | 8/18 (44%) | 14/18 (78%) | 14/18 (78%) | 4/18 (22%) | 0/10 (0%) | 17/18 (94%) |
| Average <sup>1</sup> | 25/116 (22%) | 90/120 (75%) | 40/116 (34%) | 24/116 (21%) | 10/121 (8%) | 66/118 (56%) |

| Region #1/ Pathogen | Spiroplasma | Propionigenium | RLB | Vibrio | AHPND | Microsporidia |
| --- | --- | --- | --- | --- | --- | --- |
| PL | 2/7 (29%) | 4/6 (67%) | 0/6 (0%) | 0/6 (0%) | 0/6 (0%) | 2/6 (33%) |
| Farm | 14/37 (38%) | 27/42 (64%) | 6/38 (16%) | 9/38 (24%) | 9/42 (21%) | 22/40 (55%) |
| Broodstock | 0/24 (0%) | 24/24 (100%) | 14/24 (58%) | 9/24 (38%) | 1/24 (4%) | 8/24 (33%) |
| Wild Broodstock | 0/12 (0%) | 12/12 (100%) | 0/12 (0%) | 0/12 (0%) | 0/12 (0%) | 9/12 (75%) |
| Region #2 |  |  |  |  |  |  |
| PL | 1/1 (100%) | 0/1 (0%) | 1/1 (100%) | 0/1 (0%) | 0/1 (0%) | 0/1 (0%) |
| Farm | 0/17 (0%) | 9/17 (53%) | 5/17 (29%) | 2/17 (12%) | 0/17 (0%) | 8/17 (47%) |
| Region #3 |  |  |  |  |  |  |
| Farm | 8/18 (44%) | 14/18 (78%) | 14/18 (78%) | 4/18 (22%) | 0/10 (0%) | 17/18 (94%) |
| Average <sup>1</sup> | 25/116 (22%) | 90/120 (75%) | 40/116 (34%) | 24/116 (21%) | 10/121 (8%) | 66/118 (56%) |

\* If different methods were available highest value was taken as positive.
\*\* There are multiple geographical variants of IHHNV, some of which are not distinguishable by all the available methods for IHHNV (Lightner, 2011). Several primers sets have shown utility in routine use. Among these are 70012F/77353R, 389F/R, 392F/R, MG831F/R and IHHNV 309F/R, when using a panel of primers, if some but not all primers amplify, this pattern is a clear indication that the PCR primers are measuring an endogenous viral element (EVE) and not complete virus (Brock et al. 2022).
\*\*\* Zhang et al. (2014).

**Table 11.** Comparison of prevalence/average of copies of *Propionigenium* in *P. vannamei* in samples from 3 different regions of Latin America.

|  | PG16S-F/R<br>(Munkongwongsiri, et al. 2022) |
| --- | --- |
| **Region #1** |  |
| PL | 4/6 (67%)<br>8.9 x 10<sup>1</sup> copies*<br>(0 to 2.5 x 10<sup>2</sup>) ** |
| Farm | 20/26 (75%)<br>1.27 x 10<sup>5</sup> copies<br>(0 to 3.2 x 10<sup>6</sup>) ** |
| Broodstock | 24/24 (100%)<br>1.49 x 10<sup>5</sup> copies<br>(1.3 x 10<sup>1</sup> to 2.4 x 10<sup>6</sup>) ** |
| Wild Broodstock | 12/12 (100%)<br>7.6 x 10<sup>1</sup> copies<br>(1.3 x 10<sup>1</sup> to 2 x 10<sup>2</sup>) ** |
| **Region #2** |  |
| PL | 0/1 (0%) |
| Farm | 9/17 (53%)<br>7.5 x 10<sup>4</sup> copies<br>(0 to 1.3 x 10<sup>6</sup>) ** |
| **Region #3** |  |
| Farm | 7/10 (70%)<br>1.88 x 10<sup>3</sup> copies<br>(0 to 8.3 x 10<sup>3</sup>) ** |
| Average <sup>1</sup> | 20/26 (75%)<br>1.27 x 10<sup>5</sup> copies<br>(0 to 3.2 x 10<sup>6</sup>) ** |

\*Average of copies
\*\* Range

**In Region 1**, WzSV8, *Propionigenium* and microsporidia were present in all the samples (88%, 83% and 49%, respectively, as averages of all sites), the presence in all samples including wild animals would suggest that these 3 microorganisms are endemic in *P. vannamei*. CMNV was present in all except wild animals with an average prevalence of 24%, being more abundant in post larvae samples (50%). *Spiroplasma* was found only at the hatchery and farm level samples (29% and 38% respectively). RLB, *Vibrio* and AHPND were found in only farm and broodstock samples (37%, 31% and 13%, respectively, as an average in two sites in region 1. DHPV and WSSV were found in farm animals (19% and 23% respectively), and DHPV (33%) in wild animals and WSSV (13%) in broodstock.

IHHNV infectious form and its endogenous non-viral form was only found in farm animals (15% and 23% respectively). By using the primers that target the EHP SWP gene as the most selective primer set, only 1 of 42 samples was positive (Table 9).

**In Region 2**, WzSV8 and RLB were present in both larvae (only one sample) and farm samples (100% and 29% respectively). Spiroplasma was detected only in a single sample of larvae. For farm animals the detection prevalence was 53%, 47%, 18%, 12% and 6%, respectively, for *Propionigenium*, non EHP-microsporidia, DHPV, *Vibrio* and IMNV (only found in one of 17 samples).

**In Region 3** only animals from farms were received. Major microorganisms detected were microsporidia, WzSV8, RLB, *Propionigenium* and *Spiroplasma* at relatively high prevalence of 94%, 83%,78%, 78% and 44%, respectively, while DHPV, DIV1 (5 of 18 animals were found positive), *Vibrio*, WSSV and EHP were detected a much lower prevalence of 28%, 28%, 22%, 6% and 6%, respectively (tables 9 and 10). The high prevalence of EHP (56%) (table 9) using the Tourtip method (Tourtip et al. 2009) vs the other 4 methods (0-6%) is clear evidence that there was cross reaction with other microsporidia (Jaroenlak et al. 2016). It is interesting to note the high prevalence of *Spiroplasma* using primers other than those for *S. penaei*. If this is correct, it could mean that other species of *Spiroplasma* are already infecting *P. vannamei* in the region (Ding et al. 2007, Liang et al. 2010).

### 3.3. Miscellaneous PCR findings from broodstock specimens

Ovaries of four broodstock were collected for PCR analysis. We found that all of them were positive for WzSV8 and *Propionigenium* (copy number ranged from 36 to 1,000/sample). The detection of WzSV8 in the gonads by PCR coincides with the detection by histological sections and raises concern whether WsSV8 can affect these two completely different organs, but it also suggests that this virus may be transmitted both horizontally and vertically.

### 3.4. Histology results by frequency of lesions

It is interesting to note that viruses were the dominant microorganisms with pathogenic potential in all 3 regions. If we exclude any other pathological alteration besides viral inclusion bodies, 80% of all samples analyzed (124 samples) presented at least one type of viral inclusion (VIN), and 18% at least 2 types of VIN.

In all regions WzSV8 was the most common virus observed, with the HP as is main target organ. Seventy five percent VIN were due to WzSV8. WzSV8 VIN were found in all tubule’s epithelial types except F-cells (**Pic. 1**) but focusing on E-cells facilitates rapid screening because of their location on the outer rim of the HP and because they rarely contain vacuoles. Lightner double inclusions (LDI) (Srisala et al. 2022, 2023) can be considered pathognomonic for WzSV8 because of their unique morphology, of an additional (usually smaller) eosinophilic inclusion adjacent to the single, circular basophilic, inclusions in vacuoles (**Pic. 1**). In some specimens rounded up, sloughed WzSV8 infected cells were noted in the lumen of tubules. Interestingly, WzSV8 VIN including both basophilic and LDI were also found frequently in ovaries of some specimens (PCR positive for WzSV8), (**Pic. 2**), at both developing and mature stages. In addition, WzSV8 VIN were occasionally found in the anterior midgut caecum (**Pic. 2**).

DHPV was the second most common virus lesion encountered with 18% of shrimp examined and having VIN indicative of infection. DHPV VIN are characterized as basophilic VIN that are intact bodies (viral inclusion bodies or VIB) that can sometimes be seen, still intact, after being sloughed into the hepatopancreatic tubule lumen. During their developmental expansion in the nuclei of HP tubule epithelial cells or occasional the anterior midgut caecum (AMC), the chromatin is marginated, and the intervening nucleolus is pushed aside to sometimes resemble a “bracket” to the expanding VIB. An example is shown here in the AMC (**Pic. 3 and 4**). DHPV had a higher tropism for the AMC based on the frequency of tissue positives for VIB, namely, 87% of the time in the anterior caecum, 26% in the hepatopancreas and 4% in the posterior caecum. This contrasts with Asia where DHPV is very rarely reported from *P. vannamei* (mostly reported from *P. monodon* and only in its HP). This suggests a possible difference in strains of DHPV in Asia and the Americas.

From all samples, 48% had either nematodes or gregarines or both. Larval nematodes (**Pic. 5 and 6**) were encysted in the wall of the foregut near the junction with the midgut. Gregarine trophozoites were within the lumen of the anterior midgut caecum and midgut intestine whilst gametocytes were within the lumen of the posterior midgut caecum.

For the three regions 26% of all shrimp analyzed had lymphoid organ spheroids and more than 50% of the animals had pathological changes and/or diagnostic changes consistent with viral infection (WzSV8, HPV/DHPV, BP) in the HP. Three percent of shrimp had LOS but no indication of viral infection or host inflammatory responses in the HP (**Pic. 7**).

Histopathological alterations of the hepatopancreas were characterized by acute to chronic tubule epithelial necrosis and sloughing with host inflammatory responses around and into the affected tubules. This sometimes-included melanization as well as large tubule distention and epithelial compression (atrophy-collapsed) along the basement membrane of the affected tubules. There was scant to absent eosinophilic stain uptake within the sinusoids of the areas of tubules lacking hemocytic cellular infiltration (**Pics. 8-10**).

The pathologies observed were consistent with acute to chronic toxicity of the hepatopancreas (due to Pir A/B exotoxins) as well as intratubular microbial infections by selected microbial agents (hepatopancreatic vibriosis and intracellular bacteria) (**Pics 8-10**).

Clusters of microsporidian spores were mainly observed within striated muscle sarcomeres although examples were also noted within HP tubules and cells of the interstitium (**Pic. 11**).

### 3.5. Histology results by region

**Region 1** had the highest range of identified pathogens, excluding microsporidia and encysted nematodes. The majority were within the HP/mid-gut with the exceptions of WSSV (not in the HP) and WzSV8 in the gonads (**Pics. 1-10**). WzSV8 prevalence ranged from 5% in larvae to 100% in wild animals. WzSV8 infected cells and LO spheroids were the most frequent histological anomalies: average of all samples 49% and 43% at low grade (scored at 1.3 to 1.5), respectively.

Although variable between individual shrimp reactive cells, nodules within LOS contained cytoplasmic vacuoles, karyorrhectic to pyknotic nuclei and/or suspected small RNA virus inclusions. The histopathology of wild animals (average weight 43 g) indicated endemic status of WzSV8 in the native population of *P. vannamei* in the region (**Table 12**). In farm and broodstock shrimp the prevalence of WzSV8 was 44% and 47%, respectively. DHPV prevalence in region 1 samples was 12%, 1% and 30%, for farm, broodstock and wild shrimp. Average lesions in the hepatopancreas and intestine both at grade 1.5 were noted at 12% and 1.0%, respectively.

**Table 12:** Histological average of lesions in the 3 regions.

| Region # |  |  | Collapsed<br>Hepatopancreas <sup>1</sup> |  | Intestine |  | Muscle |  | Exoskeleton |  | LO |  |
| --- | --- | --- | --- | --- | --- | --- | --- | --- | --- | --- | --- | --- |
|  | Ave <sup>3</sup>(g) | n <sup>4</sup> | % <sup>5</sup> | grade <sup>6</sup> | % <sup>5</sup> | grade<sup>6</sup> | % <sup>5</sup> | % <sup>5</sup> | % <sup>5</sup> | grade<sup>6</sup> | % <sup>5</sup> | grade<sup>6</sup> |
| Region #1 |  |  |  |  |  |  |  |  |  |  |  |  |
| Larvae | 0.01 | 20 | - | - | - | - | - | - | - | - | - | - |
| Farm | 12.7 | 200 | 12.8 | 1.9 | 2.0 | 2.0 | - | - | 2.2 | 0.3 | 50.8 | 1.9 |
| Broodstock | 39.9 | 108 | 22.2 | 1.4 | 0.3 | 1.0 | 1.6 | 0.8 | 1.6 | 0.2 | 59.3 | 1.7 |
| Wild Broodstock | 42.9 | 10 | 10.0 | 1.1 | - | - | - | - | - | - | 60.0 | 2.5 |
| Region #2 |  |  |  |  |  |  |  |  |  |  |  |  |
| PL | ND<sup>7</sup> | 10 | - | - |  |  | - | - | - | - | - | - |
| Farm | 9.1 | 65 | 3.8 | 0.8 | 6.7 | 1.3 | 0.7 | 1.0 | 10.0 | 1.3 | 6.0 | 1.5 |
| Region #3 |  |  |  |  |  |  |  |  |  |  |  |  |
| Farm | 16.8 | 59 | 11.1 | 1.1 | - | - | - | - | - | - | 7.6 | 1.0 |

| Region # |  |  | WSSV<sup>8</sup> |  | HPV<sup>8</sup> |  | WzSV<sup>8</sup> |  | Microspor |  | Gregar/Nema<sup>2</sup> |  |
| --- | --- | --- | --- | --- | --- | --- | --- | --- | --- | --- | --- | --- |
|  | Ave <sup>3</sup>(g) | n <sup>4</sup> | % <sup>5</sup> | grade <sup>6</sup> | % <sup>5</sup> | grade<sup>6</sup> | % <sup>5</sup> | % <sup>5</sup> | % <sup>5</sup> | grade<sup>6</sup> | % <sup>5</sup> | grade<sup>6</sup> |
| Region #1 |  |  |  |  |  |  |  |  |  |  |  |  |
| Larvae | - | - | - | - | 5.0 | 1.0 | - | - | - | - | - | - |
| Farm | 4.0 | 3.7 | 11.5 | 1.9 | 43.5 | 1.7 | 2.5 | 3.0 | 25.0 | 1.2 | 4.0 | 3.7 |
| Broodstock | 2.1 | 3.3 | 0.7 | 1.7 | 46.6 | 1.6 | - | - | 24.0 | 1.3 | 2.1 | 3.3 |
| Wild Broodstock | - | - | 30.0 | 2.5 | 100 | 1.0 | - | - | 90 | 1.1 | - | - |
| Region #2 |  |  |  |  |  |  |  |  |  |  |  |  |
| PL | - | - | - | - | 26.0 | 1.0 | - | - | - | - | - | - |
| Farm | - | - | - | - | 44.5 | 1.1 | - | - | -- | -- | - | - |
| Region #3 |  |  |  |  |  |  |  |  |  |  |  |  |
| Farm | - | - | 4.0 | 1.0 | 31.1 | 1.1 | - | - | 40.4 | 1.1 | - | - |

1. Average hepatopancreas abnormalities (Cell sloughing, hemocytic – melanized and necrotic tubules, atrophied/ destructed tubules, hemocytic enteritis) 2. Sum of gregarines and nematodes. 3. Average weight (g). 4. Total number of animals analyzed. 5. Percentage of prevalence. 6. Average of a grading system of severity was adopted from Lightner (1996) and simplify as follow: 0= no lesions, 1=lesions or infection present in <25% of area or organ or tissue section, 2=lesions or infection present in 25-50% of area or organ or tissue section, 3=lesions or infection present in 50-75% of area or organ or tissue section, 4=lesions or infection present in >75% of area or organ or tissue section (Lightner, 1996). 7. ND Not determined. 8. As inclusion bodies.

WSSV was detected only in farms and broodstock at a prevalence of 4.0 and 2.1%, but high grade 3.7 and 3.3, respectively. BP was found only in few farms and broodstock animals at a very low % (<2%, not included in the table). Microsporidia (2.5%) were found only in few farm animals at high grade 3 (**Pic.11**). Gregarines plus nematodes were noted in 25% of the farm and broodstock shrimp and 90% in wild shrimp.

**Region 2** had the lowest number of pathogens present. Like Region 1 WzSV8 VIN was predominant present in 26% of the larvae samples at low grade 1. In farm animals the prevalence of WzSV8 averaged 45% at grade 1 (**Pic. 12**). Systemic inflammatory responses were noted in 11% of the specimens (**Pic. 13**). These changes were hemocytic aggregations in hemocytic nodules, some of which were melanized, in the lymphoid organ, heart, sinusoids of the hepatopancreas, hematopoietic tissue and connective tissue of abdominal muscle and the stomach (**Pic. 14 and 15**). The prevalence of specimens showing LOS was 6%.

**Region 3** virus prevalence was 31% and 4% for WzSV8 and DHPV, respectively (**Pic. 16 and 17**). The other viruses were not detected by histology in this region. Forty percent of the specimens had gregarines plus encysted nematode larvae associated with the digestive track (**Pic. 18**). Hepatopancreas necrotic tubules and prominent host inflammatory responses in the surrounding sinusoids were found. Eleven % prevalence of abnormal HP was found in this region (**Pic. 19**).

### 3.6. Concurrence of histological changes

It is interesting to note that viral presence was widespread in all 3 regions. In Region 1, excluding any other pathological anomaly and only focusing on the presence of viral inclusions in farmed, broodstock or wild shrimp, respectively, revealed that 77%, 83% and 100% of the samples had at least 1 virus and that 25%, 17% and 30% had at least 2. In region, 2, 73% of the samples analyzed had at least one virus present and in region 3, 80% of the samples had one virus present and only 7% had 2.

In all regions WzSV8 VIN was predominant. While WzSV8 VIN was always found mainly in the hepatopancreas, DHPV was found 87% of the time in the anterior caecum, 26% at the hepatopancreas and 4% in the posterior caecum. This contrasted with the type of DHPV that occurs in *P. monodon* in Asia where its inclusions occur only in the HP and where DHPV is very rarely reported in farmed *P. vannamei*. These results may be the result of varietal strains of DHPV, from which at least 8 full genome sequences are available at GenBank (Srisala et al., 2021).

DHPV was the second most important viral pathogen present in regions 1 and 3. In Region 1 it was present at 12%, 1% and 30% in farms, broodstock and wild animals, respectively. While in Region 3 prevalence was 5%. In Region 1, we also found WSSV and BP in farm animals and broodstock only, with WSSV prevalence at 4.6% in farm animals and 0.5 % in broodstock, and with BP prevalence at 2.6% in farm animals and 2.5% in broodstock. While BP was found exclusively in the hepatopancreas, WSSV was found in the stomach, antennal gland, connective tissue, gills, epidermis, and gonads.

In Region 1, combined nematodes and gregarines were found in farm (26%), broodstock (28%) and wild animals (90%). In Region 3, 42% were found with the same parasites. No parasites of this kind were found in region 2. These parasites were found mostly in the anterior and posterior part of the intestine and the anterior midgut caecum.

### 3.7. Consistency between PCR and histology results

WzSV8, DHPV and WSSV were the only pathogens that were seen in histopathological samples (unequivocally identified by their viral inclusions) and at the same time detected by PCR. However, it is important to note that PCR and histology results in this report did not arise the same animals, but from random samples of the same population from the same pond. Other pathogens such as *Vibrio spp*, AHPND associated bacteria and RLB were detected in high prevalence by PCR. Unfortunately, those observations could not be related directly to the histological damages seen in the hepatopancreas or other tissues. Nor could we confirm the identity of the bacteria in the lesions because we lacked tools such as *in situ* hybridization. For this reason, we could not identify the pathogens in the tissue sections, and we simply referred these pathogens as bacteria.

## 4.0. DISCUSSION

The present study describes the histopathological lesions and agents found by PCR in *P. vannamei* samples collected from 3 different regions in Latin America. It also compares different sets of primers for a range of agents to understand possible geographical variations and their degree of specificity. We also highlighted the importance of selecting the correct set of PCR primers specific for the target agents which may vary with geographical location. Another important aspect that should be highlighted in this study is the difference between the level of coinfection between the 3 regions, which is related to the culture system or the health status of the broodstock/source of the post larvae. Region 1 is characterized by zero biosecurity, massal selection, broodstock go from the farm to the hatchery regardless of their health status. The animals from region 2 were sampled from a single farm with a super intensive close culture model with biosecurity and their broodstock are originated from a close system not from the farm. The region 3 animals come from an open farm without biosecurity, but the broodstock and larvae were SPF.

The lack of consistency found in several primers for viruses could be attributed to the genetic diversity of pathogens affected by geographical distribution of the host, environmental factors, and adaptations of the viruses to different environmental factors in different geographical areas (Safeena et al. 2012). Existing PCR detection methods can also give false positive test results due to cross reactivity of the primers with closely related organisms (Jaroenlak, et al. 2016). There is also the possibility of false positive PCR test results arising from EVE that include the sequence of the PCR target. Finally, the accuracy of the libraries obtained using universal primers depends strongly on the choice of primers (Baker et al. 2003, Klindworth et al. 2013). More studies on this subject should be carefully considered.

WzSV8 was the most common pathogen detected by PCR and by histology in all regions sampled regardless of size of the animal and/or the environment. WzSV8 was rarely found alone. Generally, WzSV8 was accompanied by lesions in the mid-gut presumed to be due to bacteria, and to a lesser prevalence of HPV/DHPV, WSSV or BP. WzSV8 was found in cultured larvae as well as in wild animals (Region 1), the later at 100% prevalence. So, it is not wrong to assume that this virus is endemic to this region and already widespread in the world.

Srisala et al. (2022, 2023) were the first to identify and characterize WzSV8 lesions in hematoxylin and eosin (H&E) stained tissues using a light microscope. Grossly normal farmed *P. vannamei* juveniles from Asia were screened for WzSV8 by RT-PCR using primers designed from the WzSV8 RNA sequence (KX883984.1) submitted to GenBank (Li et al., 2015). Positive amplicons were used to prepare a DNA *in situ* hybridization (ISH) probe to examine the tissues of the shrimp that had given the positive RT-PCR results. Positive ISH results were seen in the hepatopancreas (HP) and were outstanding within vacuoles of the E-cells. Comparison with adjacent tissue sections stained with hematoxylin and eosin (H&E) revealed circular, deeply basophilic inclusions within vacuoles in the cytoplasm of tubule epithelial cells of the HP. Cruz-Flores et al. (2022) reported yet another type of WzSV8 from Brazil with high genome sequence similarity to the sequences of WzSV8 reported from Thailand, China, and Australia (Li *et al*. 2015, Huerlimann *et al*. 2018, Liu et al, 2021). Based on recent changes in viral taxonomy, they named it Penaeus vannamei solinvivirus (PvSV). In contrast to the study of Srisala et al. (2023) above, the ISH results shown by Cruz-Flores et al. (2022) were seen only in morphologically normal HP tubule cell nuclei. They reported no ISH positive cytoplasmic inclusions in the HP nor in the ovary. It is important to mention that a structure similar to LDI in the hepatopancreatic cells has been reported by Owens and co-authors while describing the presence of Lymphoid Parvo-like virus in *P. monodon* (Owens et al. 1991, Owens, 2023). In summary, it is not yet proven that WzSV8 and related viruses are shrimp pathogens that result in a disease of shrimp. In the present study, WzSV8 was mainly found in the hepatopancreas, but also found in the anterior caecum and sometimes in the gonads.

Both WzSV8/PvSV and DHPV have an affinity for the hepatopancreas. Cruz-Flores et al. (2022). suggested that this makes the HP more susceptible to co-infection with enteric diseases such as bacteria and or EHP. Cruz-Flores et al. (2022) concluded that the coinfection PvSV with infectious myonecrosis virus (IMNV) was the likely cause for the unusual mortalities that are currently affecting Brazilian shrimp farming. However, no histology or other evidence of the latter are available from Brazil. Nor have River’s Postulates been demonstrated.

In this study the midgut histopathology results are compatible with SHPN, chronic AHPND and or RLB/NHP. Unfortunately, we cannot relate these names to any of the infections/alterations of the digestive system seen since we lacked capability for *in situ* hybridization. Therefore, we cannot confirm that the pathogens in the lesions correspond to those identified by PCR using specimens of different shrimp from the same source. However, over 55% of the PCR samples tested positive for *Vibrio*, RLB and AHPND, and 12% of the histopathological lesions were in the hepatopancreas and intestine, (average after combining samples in the 3 regions). Therefore, it would not be wrong to suggest that any of these bacterial pathogens could be associated with these lesions. These findings are also consistent with those described by Morales-Covarrubias et al. (2018) and Aranguren Caro et al. (2020), who found that *Vibrio* species associated disease are predominant in the region.

The name decapod hepanhamaparvovirus (DHPV) has recently replaced the name HPV (Pénzes et al. 2020). It was one of the most common pathogens detected in the present study. Average prevalence of HPV/DHPV by PCR and histology was 30% and 18%, respectively. Shrimp affected by the DHPV usually show nonspecific gross signs, but if they become infected by other pathogens, the combination of agents can result in atrophy of the hepatopancreas, anorexia, poor growth rate, reduced preening activities, increased tendency to surface, and gill fouling by epicommensal organisms (Flegel et al. 1999., Dhar et al.2014).

IHHNV and DHPV belong to the same family *Parvoviridae* and sub-family Densovirinae but in different genera. IHHNV (genus *Penstyldenosvirus*) is a small, naked, ssDNA virus that first emerged in mid-1981 and could cause a condition described as ‘runt deformity syndrome’ (RDS) in *P. vannamei* in which affected shrimp display cuticular deformities and slow growth (Kalagayan et al. 1991). Two genotypes of IHHNV have been shown to be integrated into the host genomic DNA and are not associated with histological lesions typical of IHHNV infection and experimental transmission studies suggest that they are not infectious for *P. monodon* or *P. vannamei* (Tang & Lightner 2006). In two studies (Huerlimann et al, 2022; Taengchaiyaphum et al., 2022) revealed that the sequences reported by Tang and Lightner (2006) are assembly artifacts that arose from an ancient form of IHHNV that exists in scrambled genome fragments called endogenous viral elements (EVE) that occur in clusters in the *P. monodon* shrimp genome. The study by Taengchaiyaphum et al. (2022) revealed that a separate scrambled cluster of genome fragments from a currently existing type of IHHNV also occurred in the genome of the same shrimp specimen and that it could give false positive PCR test results using methods for IHHNV detection recommended by the World Organization for Animal Health (WOAH) (Anonymous, 2017). The existence of EVE arising from IHHNV, and other viruses reveals that caution is required, especially when reporting a virus from a new location for the first time (Saksmerprome et al., 2022; Alday-Sanz et al., 2020),

The level of occurrence of IHHNV in the PCR analysis was very low and completely absent during the histopathology analysis. IHHNV was found only in farm samples from region 1. Within these samples 24% were associated with EVE while 16% might have arisen from infectious IHHNV except for absence of pathognomonic CAI lesions by histological analysis (albeit in different specimens from the same source. These findings are not rare. Rai et al. (2009) reported that non-infectious IHHNV has integrated into the genome of about one third of *P. monodon* tested. Interestingly, they also found that 22.8% of post larvae and PL and 10.5% adults had both infectious IHHNV and the EVE form (Rai et al. 2009). Another publication (Saksmerprome et al., 2011) also revealed that 40 *P. monodon* specimens in Thailand gave positive test results with a WOAH-recommended method, but that only 3 were infected with IHHNV. Brock et al. (2013) also reported that IHHNV sequences in the shrimp genome (now called EVE) could give false positive PCR test results for infectious IHHNV using WOAH (called OIE at the time) methods.

It is important realize when testing samples using multiple PCR primer sets of equal sensitivity and covering different regions of a virus genome that all the pairs must give the expected amplicons. Failure to do so is a strong indication that the primer sets giving amplicons are arising from EVE and not from the complete virus genome (Brock et al. 2023). Furthermore, IHHNV replication results in anatomic alterations within target cell nuclei visible with H&E staining or gene probe methods. We did not find any characteristic tissue alterations or Cowdry type A inclusion bodies as histological indicators of this virus. Reports of this virus causing pathology in shrimp are now very rare (Aranguren Caro et al. 2022; Romero, 2022) such that we agree with Brock et al. (2023) that IHHNV should no longer be a virus listed for shrimp by WOAH (Brock et al. 2023).

Both NHP and RLB are diseases of penaeid shrimp caused by a gram-negative intracellular rickettsia-like bacteria that target the tubular epithelial cells of the HP. RLB were very common in region 1 and 3 by PCR detection. In contrast, NHPB was almost undetected. This is another example of how selection of primers can give misleading results. We also found that the universal primer for rickettsia used in region 3 was more sensitive than the nested method designed for RLB in penaeids. It should also be considered for future studies. Unfortunately, as explained before we cannot correlate any of this PCR with histopathological lesions because we did not test using specific tools for confirmation. We did not find any histological lesion caused by *Spiroplasma*, but the detection by PCR of *Spiroplasma penaei* and other strains calls for more attention in future studies of this group.

The high prevalence of *Propionigenium* is interesting and should be considered in future studies. This strictly anaerobic Gram-negative bacterium is found in marine habitats, typically in sediments. Munkongwongsiri et al. (2022) suggested that *Vibrio* and *Propionigenium* should be tested in combination with EHP as potential component causes of EHP-White Feces Syndrome in *P. vannamei*. There were no reports of white feces in our survey, but *Propionigenium* prevalence in high copy numbers could be considered as an indication of a decaying bottom environment and could be used in the future as bioindicator for risk of white feces syndrome or other problems.

Microsporidiosis is the most common and harmful disease of decapods caused by eukaryotic microbes (Sokolova et al. 2015). More than 20 species of Microsporidia belonging to 17 genera have been reported from a variety of decapod species belonging to the families Penaeidae and Caridea (Sokolova et al. 2015). Other than EHP discussed previously, the two most abundant microsporidian parasites of penaeids are *Agmasoma penaei* and *Perezia nelsoni* that have completely different life cycles, morphology, and tissue tropism. Pasharawipas and Flegel (1994) and Pasharawipas et al. (1994) designed a specific DNA probe to identify the microsporidian *Agmasoma* as the parasite infecting muscles in both *P. merguensis* and *P. monodon* in Thailand. Later Sokolova & Hawke (2016) suggested that muscle infections previously attributed to *A. penaei*, might have been due to overlooked *P. nelson* in dual infections with *A. penaei* that they believe is confined to gonads while *P. nelsoni* is confined to muscles. In the present study a 38% average PCR prevalence of microsporidia was revealed in the three regions. However, by histopathology the prevalence in muscles and the hepatopancreas was low and only seen in farm animals in region 1. Only farmed shrimp were positive. Additional study on this inconsistency is needed.

In penaeid species, the lymphoid organ has an important role in immune defense and is possibly the major phagocytic organ in penaeids (Rusaini., Owens 2010a). It is also a major site for viral degradation within lymphoid organ spheroids (LOS) (Rusaini., Owens 2010a). The extensive spheroid formation we found in all 3 regions during this study might be related to the tolerance of penaeid prawns to bacterial and viral infection. In addition, Crustaceans have a cyclic phenomenon of moulting, for development and growth that is not encountered in most other groups of aquaculture animals. Biochemical and physiological changes occur during the moult cycle, as do changes in immune components. Tidal and lunar rhythmicity also affects the physiology of some crustaceans (Rusaini., Owens 2010b). Rusaini & Owens (2010b) suggested that the elimination mechanism for spheroid cells as prawns age seems to be associated with lunar rhythms.

Generally, a pathogenic agent must be present for disease to occur. However, presence of that pathogen alone is not always sufficient to cause disease. Casadevall and Pirofski (1999) redefined a new definition of a pathogen as “A microbe capable of causing host damage; the definition can encompass classical pathogens and opportunistic pathogens; host damage can result from either direct microbial action or the host immune response”. A variety of factors can influence whether exposure to an agent will result in disease. For example: stocking density, physical and chemical factors, among others, can lead a host to become immunocompromised, and lead to disease (Armstrong,.1993; Méthot and Alizon, 2014).

In addition, it is common to attribute a viral disease to infection by a single agent. However, under natural circumstances hosts may be infected by multiple agents (coinfections), a rather frequent occurrence in penaeid shrimp. We can conclude form this survey that there is a relatively high prevalence of known shrimp pathogens in *P. vannamei* specimens from the wild, as broodstock used for PL production and as juveniles from cultivation ponds Latin America. It also revealed that many specimens had multiple infections. Thus, health outcomes may be influenced by contributions from more than a single agent, and this is often not considered in diagnostic laboratories (Diaz-Muñoz, 2017; Kumar et al. 2018). Crustaceans collected from farm and wild habitats play host to a taxonomically diverse array of DNA and RNA viruses (Bateman and Stentiford, 2017).

Multiple infections are commonly found in shrimp culture and may cause more serious consequences than infections by only one pathogen (Jang et al. 2014). For example, coinfection studies between WSSV and *Vibrio* showed that shrimp are more susceptible and that their survivability is reduced under multiple infections but may be influenced by the species of *Vibrio* bacteria (Jang et al. 2014, Pang et al. 2019). Coinfections of bacteria and a virus or a DNA virus with RNA virus in *P. vannamei* is not new. Feijo et al. (2013) reported the presence of IMNV and WSSV in Brazil. Both IMNV and WSSV don’t have the same tropism and they are highly virulent. Dewangan et al. (2022) reported multiple bacterial infections to be highly pathogenic in *L. vannamei* grow-out ponds in India. On the other hand, Flegel et al. (2004) showed that grossly normal shrimp were concurrent hosts to several viruses, and that some had significant negative correlations with shrimp size. Coinfection associated to growth retardation between EHP and DHPV was reported by Singaravel et al. (2021).

Coinfections may also result in genetic exchange between agents to generate recombinant viruses, that can influence viral evolution, disease dynamics, and eventually the fate of the host (Diaz-Muñoz, 2017; Kumar et al. 2018). Coinfections may play a pivotal role in reducing or augmenting disease severity. When individual cells are coinfected, one virus usually influences replication of the other, a phenomenon termed viral interference. Four factors that are likely to play a role in determining the type of coinfection: host ecology, host taxonomy or phylogeny, host defense mechanisms, and virus–virus interactions (Diaz-Muñoz, 2017). For example, such interactions have been reported between IHHNV and WSSV (Bonnichon et al., 2006; Melena et al., 2006) and TSV and YHV (Aranguren et al., 2012).

From a viral standpoint, persistence has benefits at different levels. A persistent infection allows virus production and assures the transmission of viral genetic material over a longer period. Because there is low or no fitness cost to the host, a persistence state could permit multiple infections (with the same or different viruses) that could be the source of new genetic variability and complexity. Since the host’s health is not significantly affected, at least in the short term, a mobile host can disseminate virus to more hosts within the same environment or to hosts in a new environment. From a host standpoint, persistence is profitable because persistently infected organisms are resistant to super infections with related viruses, a phenomenon known as viral accommodation (Flegel 2020). Organisms persistently infected with mutualistic viruses show an increased antiviral response. Mutualistic viruses can help the host by supplying new genes or through epigenetic changes of the host genome with beneficial results (Goic and Saleh, 2012). This process may take only a few years like the case of TSV or IMNV or decades as the case of WSSV. This tolerance to the pathogen does not mean resistance, so shrimp can still get infected and may develop the disease if there is a trigger (Flegel 2020).

Viral infection could also be the result of two populations of the same virus, a minority high-fitness, high-virulence phenotype, and a dominant subpopulation with a higher efficiency in progeny production in coinfected cells that interferes with replication of the virulent variants resulting in a delay of cell killing. This mirrors the evolutionary strategies of competition and colonization defined in ecology, where virulent viruses can be regarded as colonizers, because they kill the cell faster which allows them to spread faster. In turn, the interfering viruses can be regarded as competitors, because they are more efficient in exploiting the local resources which in this case are provided by individual cells (Ojosnegros et al. 2010).

NHPB: Necrotizing hepatopancreatitis bacteria
RLB: Rickettsia like bacteria
IHHNV: Infectious hypodermal and haematopoietic necrosis virus
DIV1: Decapod iridescent virus 1
HPV/DHPV: Hepatopancreatic parvovirus or hepanhamaparvovirus
MrBV: Macrobrachium rosenbergii bidnavirus
IMNV: Infectious myonecrosis virus
CMNV: Covert mortality nodavirus
PvNV: Penaeus vannamei nodavirus
MrNV: Macrobrachium rosenbergii nodavirus
YHV: Yellow head virus
EHP: *Enterocytozoon hepatopenaei*
Pv: *Penaeus vannamei*
AMC: Anterior midgut caecum

## ADKNOWLDGE

We would like to thank Dr. Victoria Alday-Sanz and Dr. Xavier Romero for the review and valuable suggestions made to this study.

---

**Pic 1:**
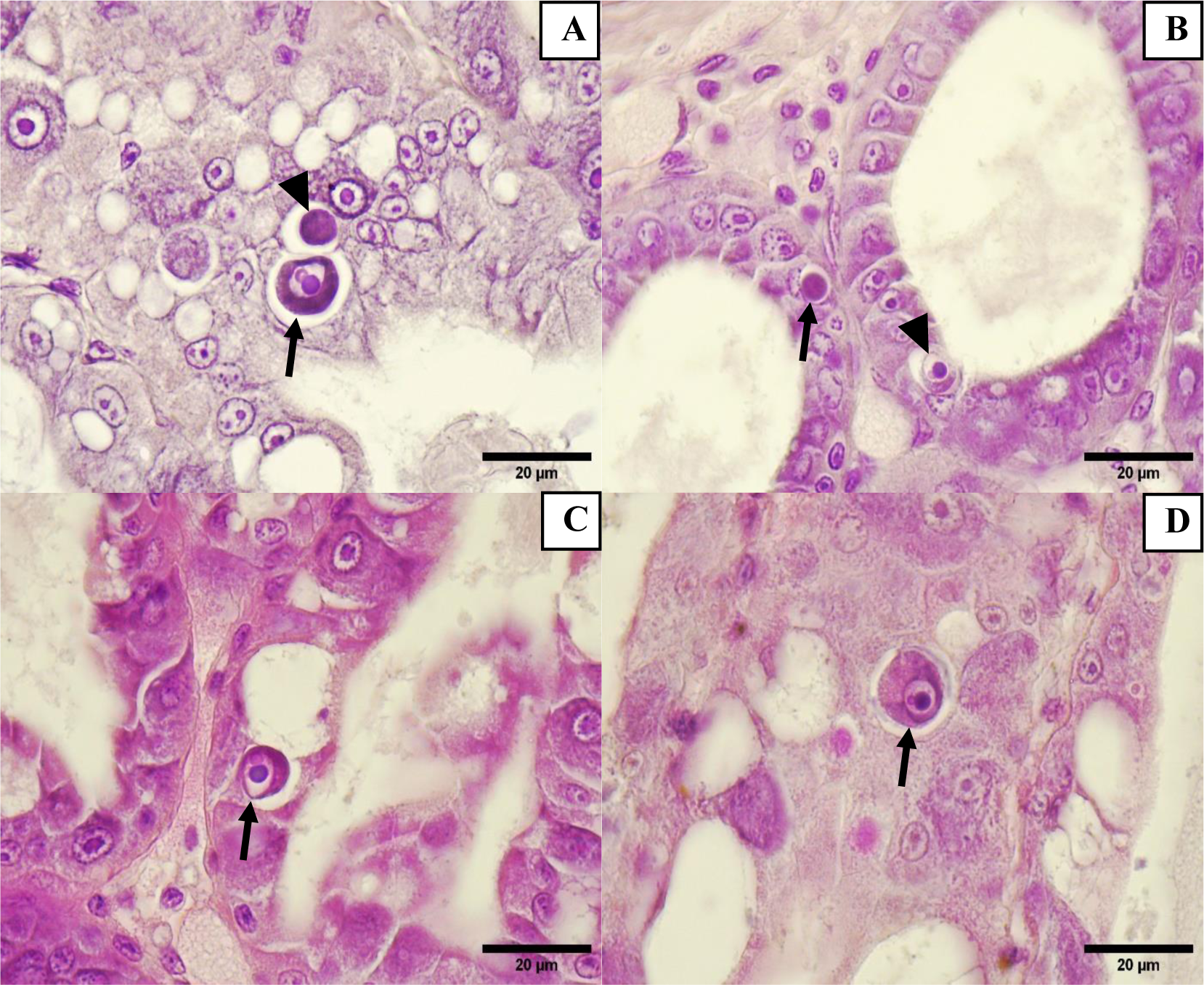
(A, B, C, D) Hepatopancreatic cells with viral inclusion body of Wenzhou Virus 8 (WzSV8), inclusion body with double Lightner (arrow) and basophilic inclusion stage (arrowhead), H&E Stain.

**Pic 2:**
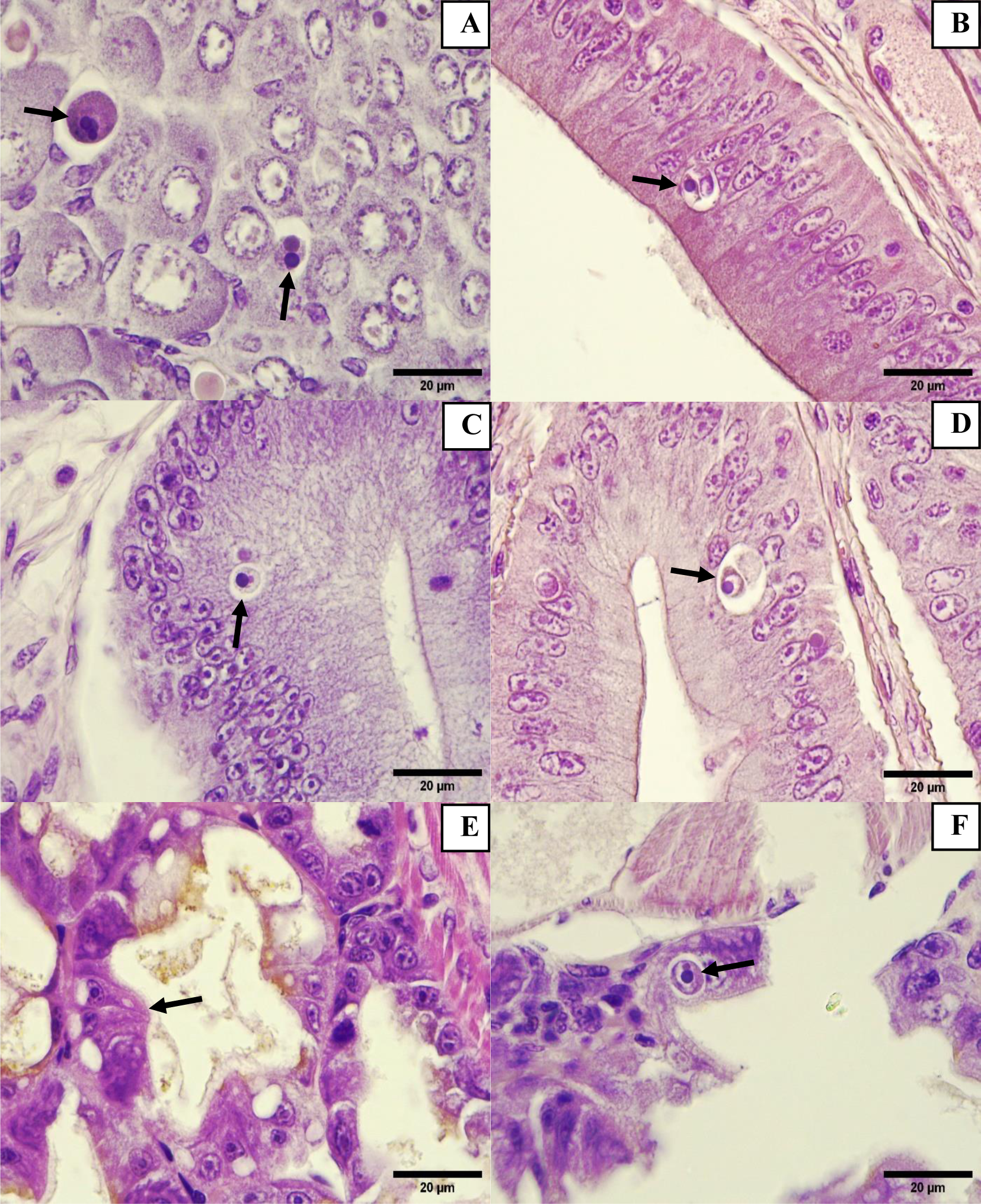
(A) Female gonad in maturation stage with WzSV8 inclusion body (arrow), 100X Magnification, H&E Stain. (B, C, D) Epithelium of anterior midgut cecum with WzSV8 inclusion bodies (arrow), H&E Stain. (E) Larvae hepatopancreas with normal tubules (arrow), 100X Magnification, H&E Stain. (F) Hepatopancreas epithelial cells with WzSV8 inclusion body (arrow), 100X Magnification.

**Pic 3:**
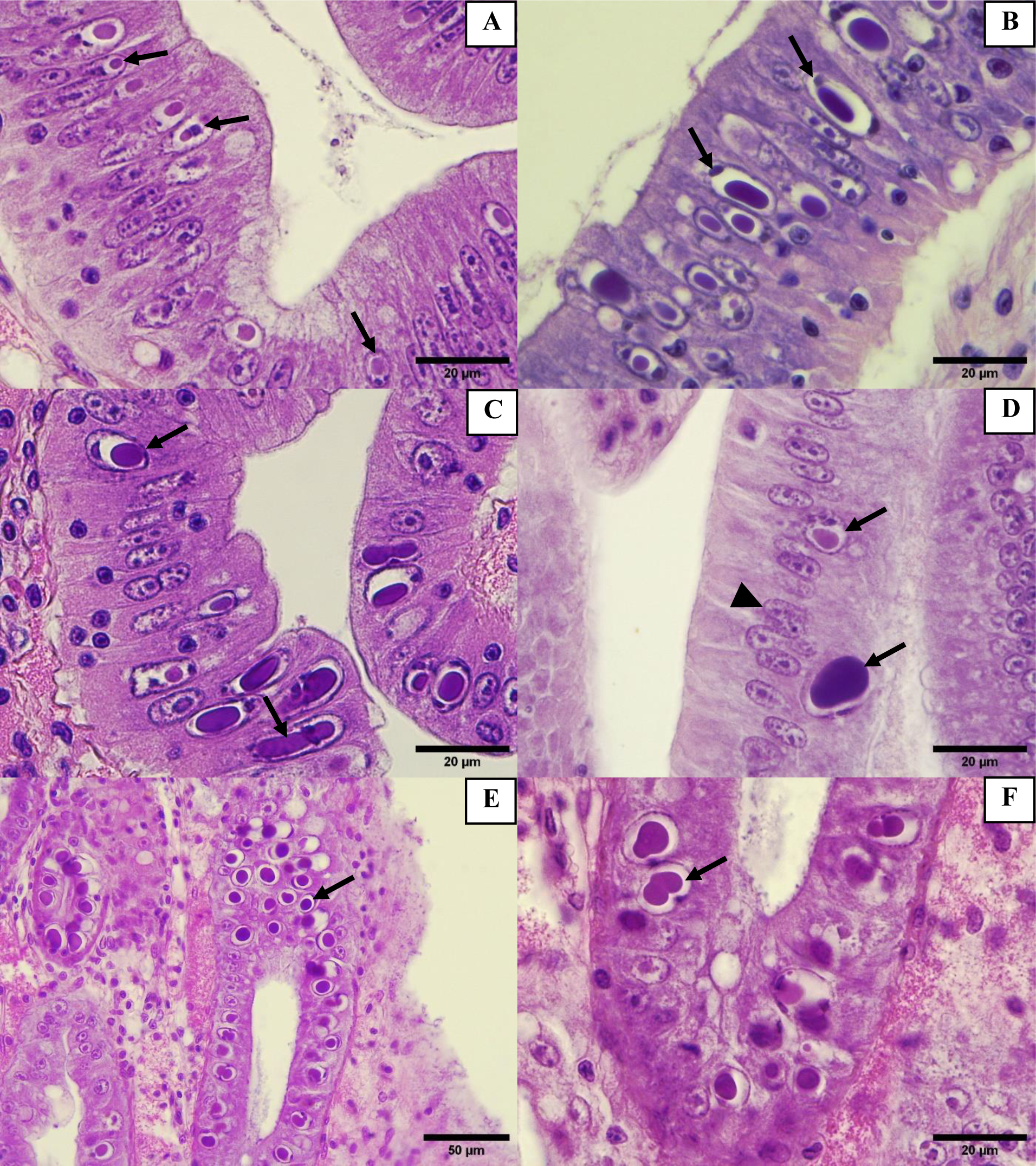
(A, B, C) Epithelium of anterior midgut cecum with DHPV inclusion bodies with compression and displacement of nucleolus (arrow), 100X Magnification, H&E Stain. (D) Epithelium of anterior midgut cecum with DHPV inclusion bodies with compression and displacement of nucleolus (arrow), in contrast, normal epithelium of anterior midgut cecum can be observed (arrowhead), 100X Magnification, H&E Stain. (E) Epithelial cells of hepatopancreas with DHPV inclusion bodies (arrow), 40X Magnification, H&E Stain. (F) Epithelial cells of hepatopancreas with DHPV inclusion bodies (arrow), 100X Magnification, H&E Stain.

**Pic 4:**
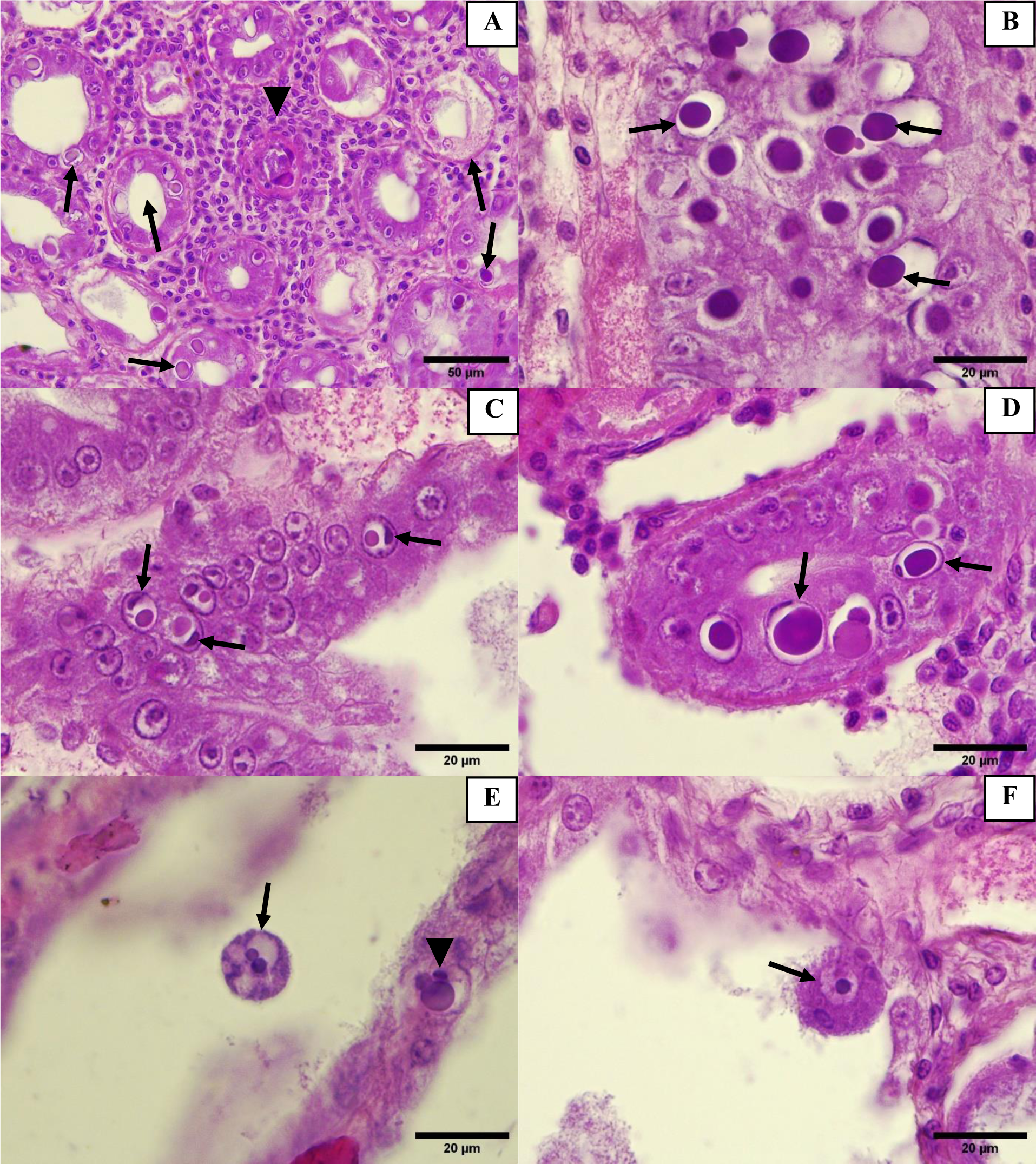
(A) Epithelial cells of hepatopancreas with DHPV inclusion bodies (arrow) and a hemocyte tubule forming (arrowhead), 40X Magnification, H&E Stain. (B, C, D) Epithelial cells of hepatopancreas with DHPV inclusion bodies with the compression and displacement of nucleolus (arrow), 100X Magnification, H&E Stain. (E) Hepatopancreas with sloughing cell, which was infected with WzSV8, the DLI was observed (arrow), the hepatopancreatic epithelial cell present an DHPV inclusion body with displacement of nucleolus (arrowhead). 100X Magnification, H&E Stain. (F) Hepatopancreas with sloughing cell, which was infected with WzSV8, the DLI was observed (arrow). 100X Magnification, H&E Stain.

**Pic 5:**
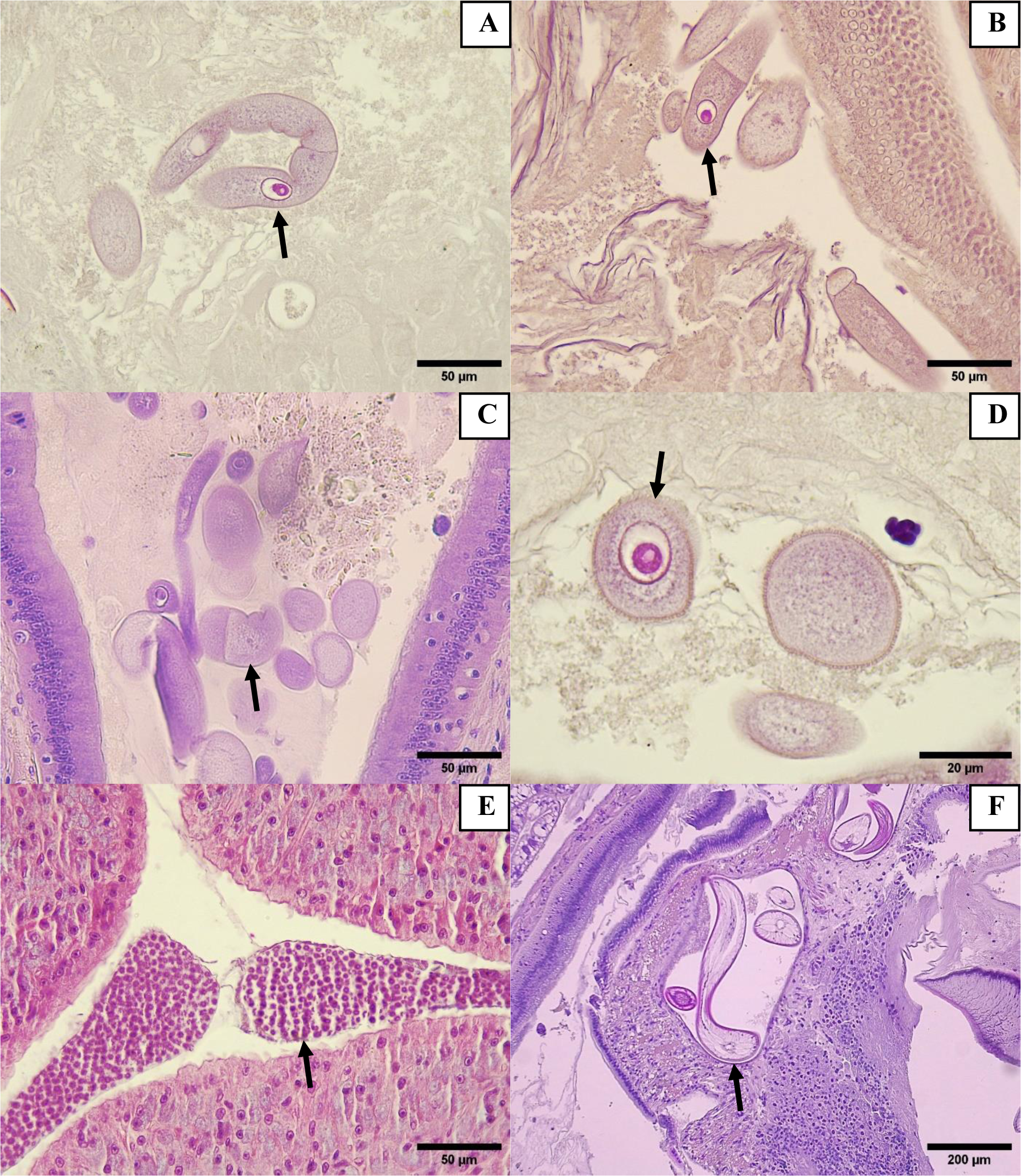
(A, B, C) Intestinal content with trophozoite stage of gregarines (arrow), Magnification 40X, H&E Stain. (D) Intestinal content with trophozoite stage of gregarines (arrow), Magnification 100X, H&E Stain. (E) Lumen of Tegumental gland of hindgut with presence of gregarines cysts (arrow), Magnification 100X, H&E Stain. (F) Connective tissue of foregut with presence of nematodes cysts (arrow), Magnification 10X, H&E Stain.

**Pic 6:**
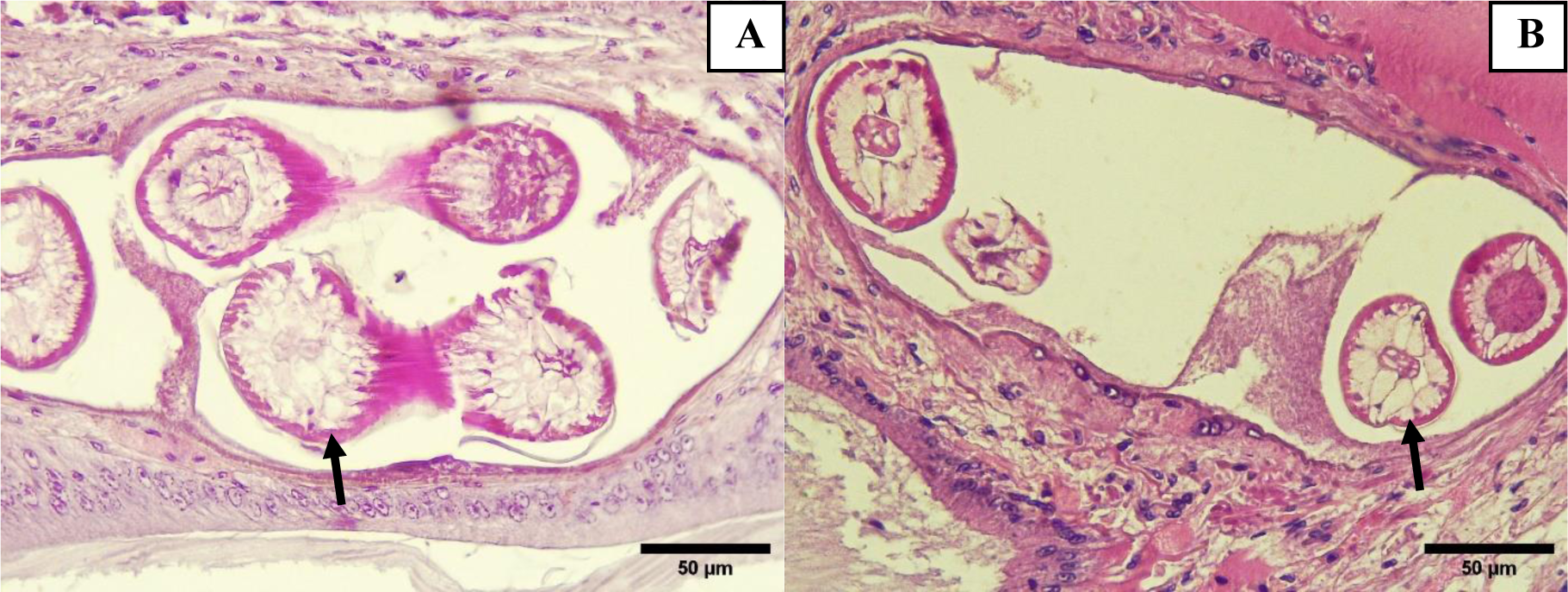
(A, B) Connective tissue of foregut with presence of nematodes cysts (arrow), Magnification 40X, H&E Stain.

**Pic 7:**
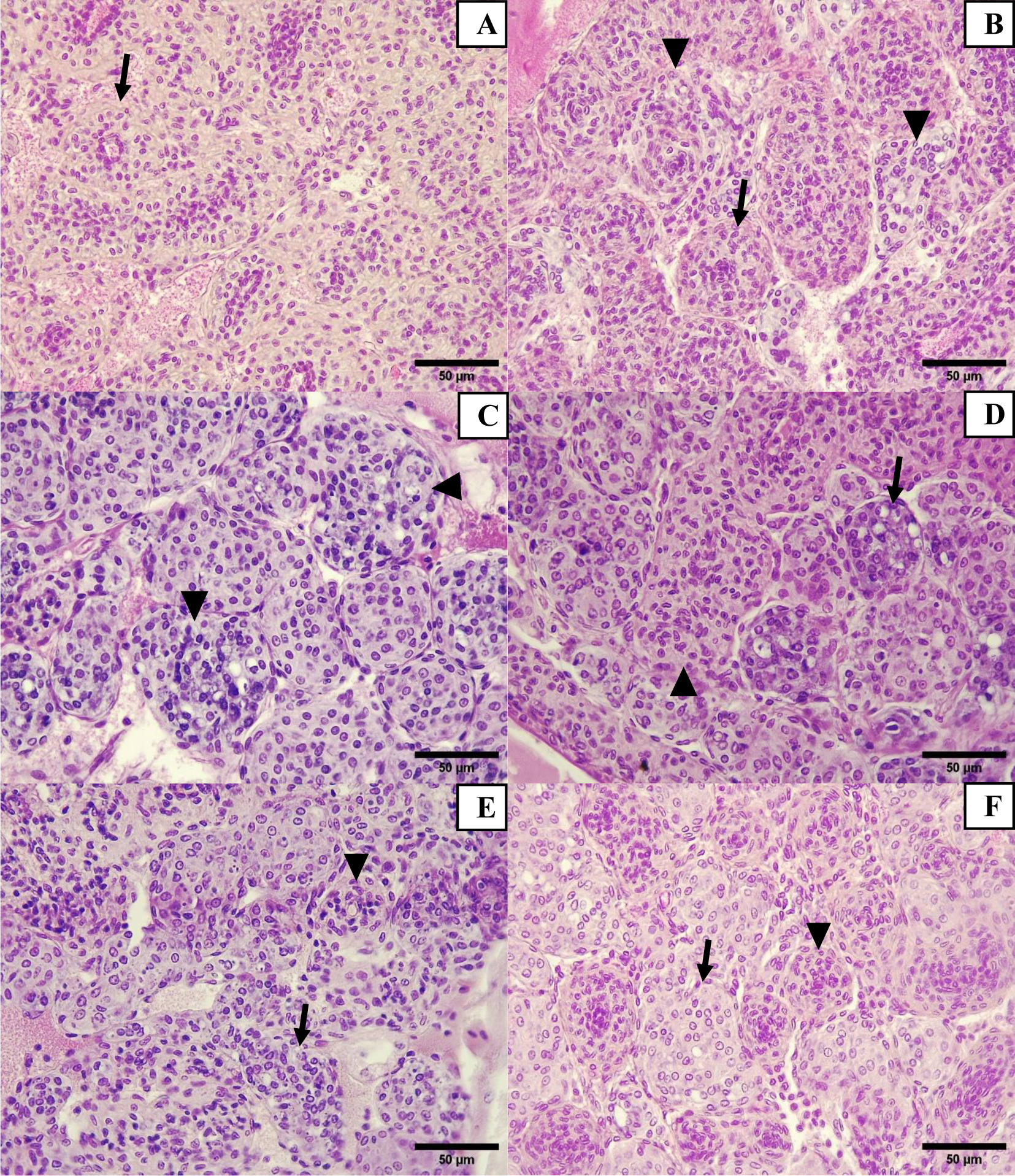
(A) Lymphoid organ: Lymphoid tubules with normal architecture (arrow), 40X Magnification, H&E Stain. (B) Normal tubule of lymphoid organ (LOS) (arrow) and Lymphoid organ spheroid (arrowhead), 40X Magnification, H&E Stain. (C, D) Lymphoid organ spheroid (LOS) with vacuoles and pyknotic nuclei (arrowhead), 40X Magnification, H&E Stain. (E, F) Lymphoid organ spheroid (LOS) with vacuoles (arrow) and normal lymphoid tubule (arrowhead). 40X Magnification, H&E Stain.

**Pic. 7.**
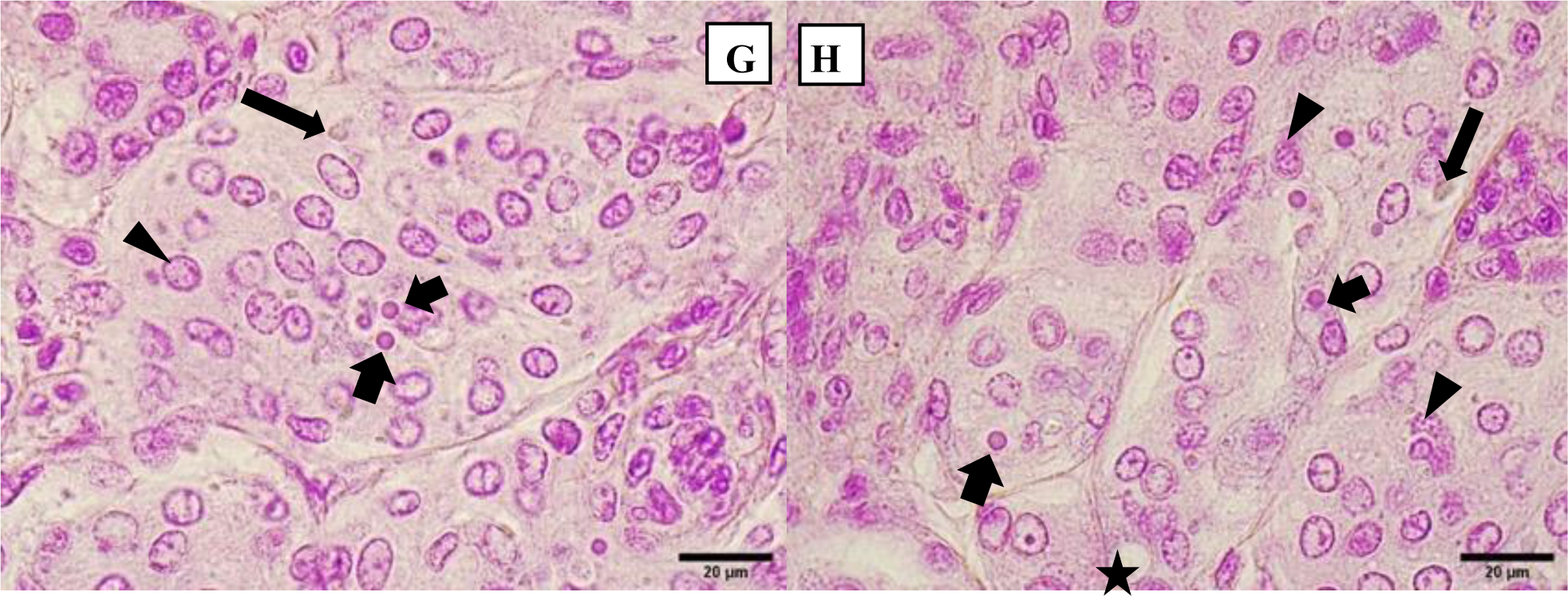
G and H, Lymphoid organ with different celular reaction: picnotic nuclei (thick arrow), cariorrexis nuclei (arrow head), viral inclusions RNA-like (Arrow) and formation of cytoplasmic vacuole (Star).

**Pic 8:**
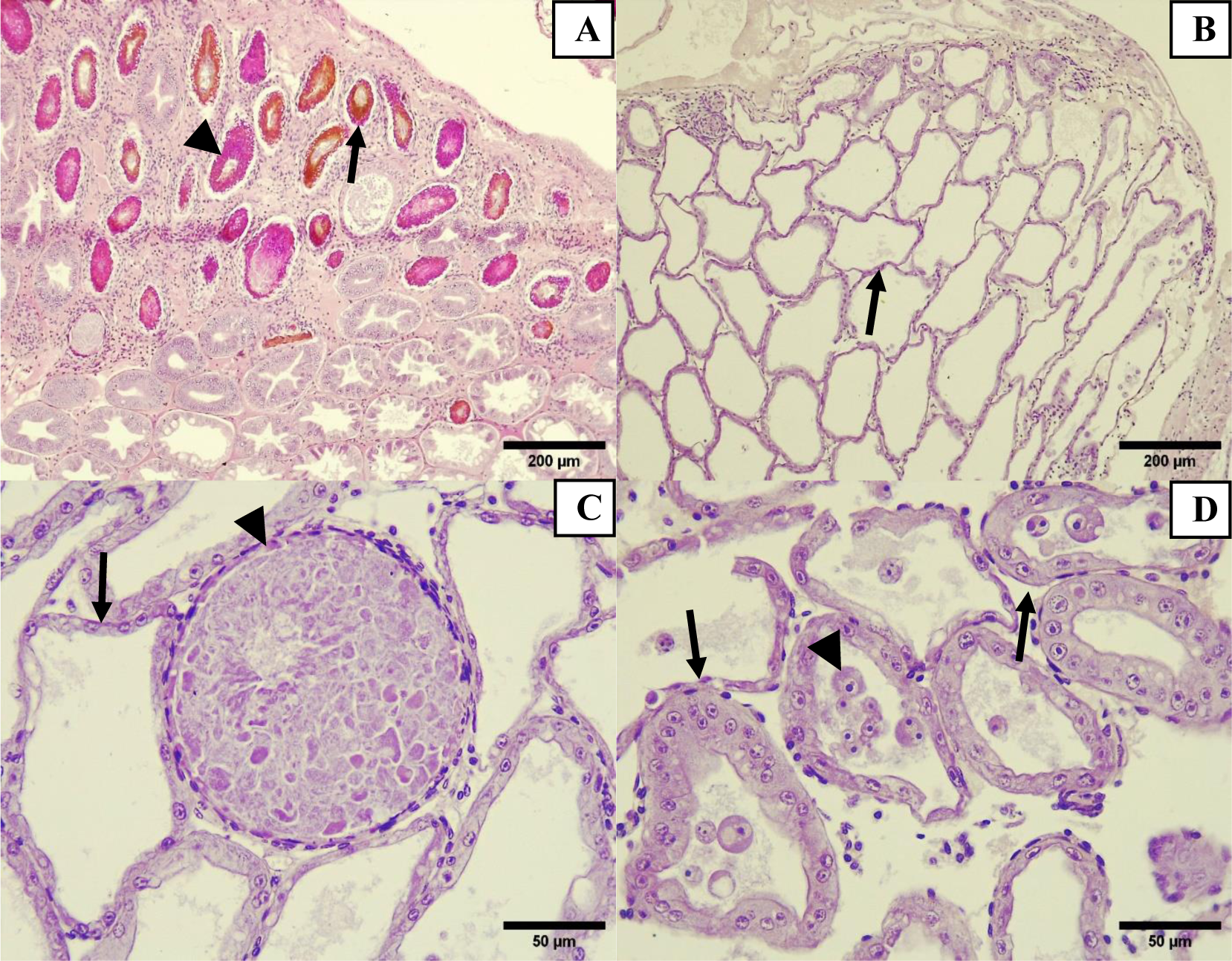
Four views of hepatopancreas from the same shrimp. (A) Large area of chronic inflammatory changes of the hepatopancreas with hemocytes infiltration and melanized, necrotic tubules (arrow and arrowhead). (Hepatopancreas in an adjacent area with acute, severe intraluminal distension, absence of intraluminal uptake of stain and atrophy (arrow) of the tubule epithelium. (C) Higher magnification view of a distended structure (a tubule or, perhaps, within a sinusoid) engorged with unrecognizable necrotic debris/bacteria. (D) Hepatopancreas histological section: Atrophied tubules without lipids or secretory vacuoles, detached WzSV8 infected epithelial cells in lumen of HP tubules and absence of eosinophilic stain uptake within the sinusoids. Note the absence of an inflammatory response H&E Stain.

**Pic 9:**
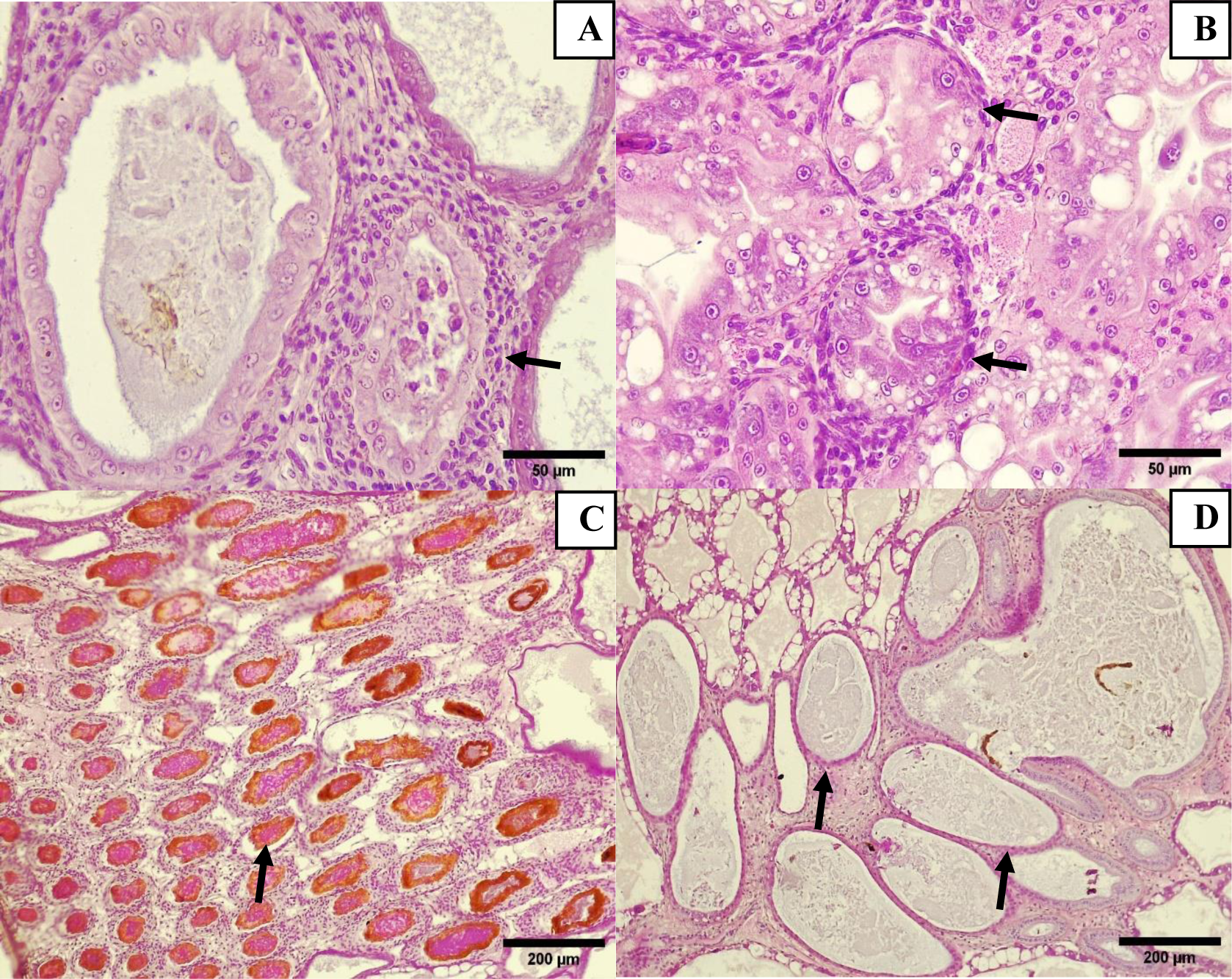
(A) Hepatopancreas histological section: Cellular debris in lumen of HP tubule surrounded with hemocytes (Arrow), 40X Magnification, H&E Stain. (B) Hepatopancreas histological section: Hemocyte surrounded of hepatopancreatic tubules (arrows), 40X Magnification, H&E Stain. (C) Hepatopancreas histological section: Destruction of tubular architecture with melanized-necrotic tubules and hemocyte infiltration (arrow), 10X Magnification. (D) Hepatopancreas histological section: Tubular architecture loss with atrophied tubules without lipids and no cell-group differentiation (arrow), 10X Magnification, H&E Stain.

**Pic 10:**
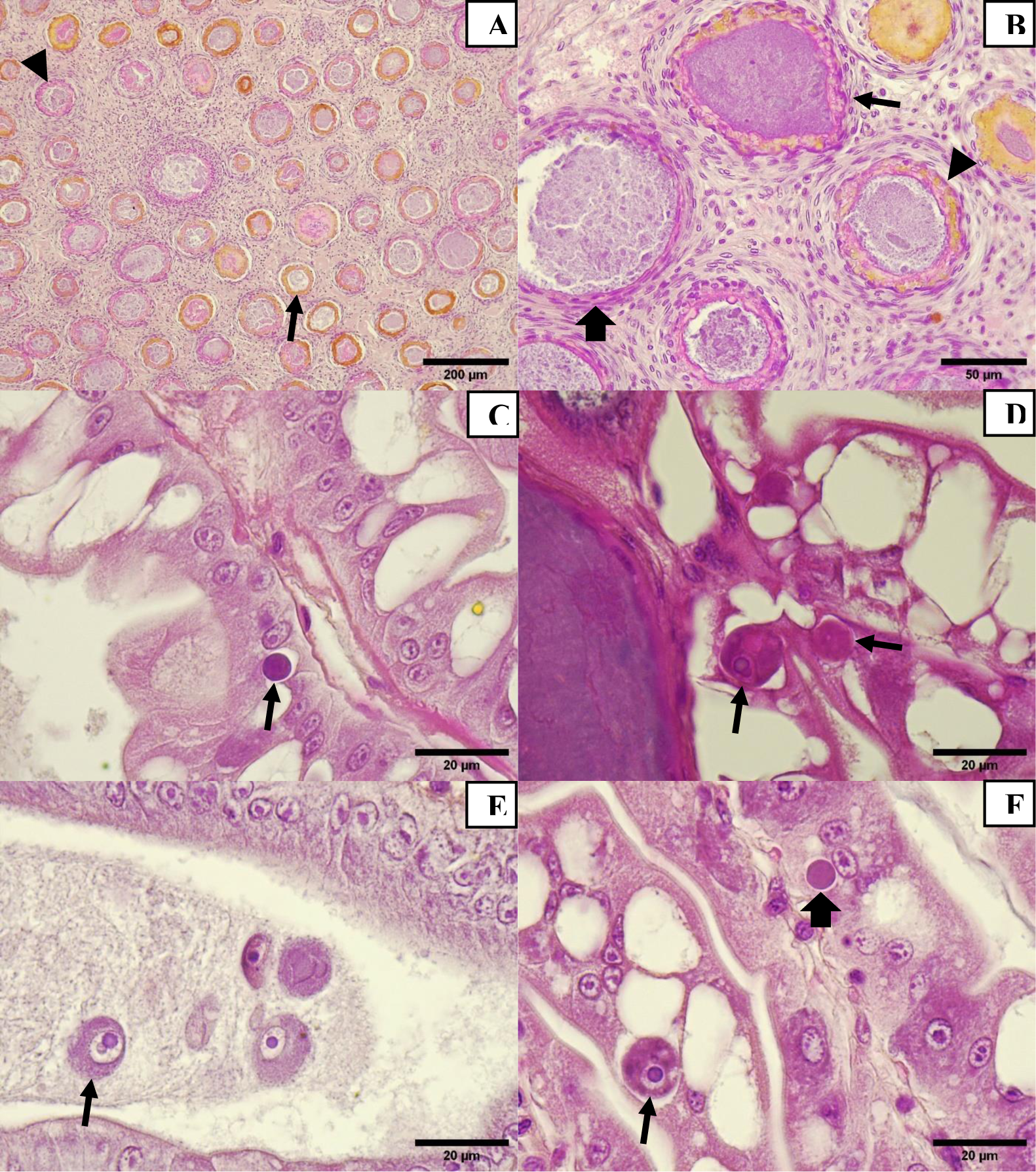
(A) Hepatopancreas histological section: Destruction of tubular architecture with necrotic tubules (arrow), melanized tubules (arrowhead), both surrounded with hemocyte infiltration, 10X Magnification, H&E Stain. (B) Hepatopancreas histological section: Tubular architecture destruction with bacterial tubule (arrow), granuloma (arrowhead) and melanized tubule (thick arrow) surrounded with hemocyte infiltration. 10X Magnification, H&E Stain. (C, D, E, F) Hepatopancreatic cells with viral inclusion body of WzSV8 we observe inclusion body with double Lightner (arrow) and basophilic inclusion stage (arrowhead), 100X Magnification, H&E Stain.

**Pic 11:**
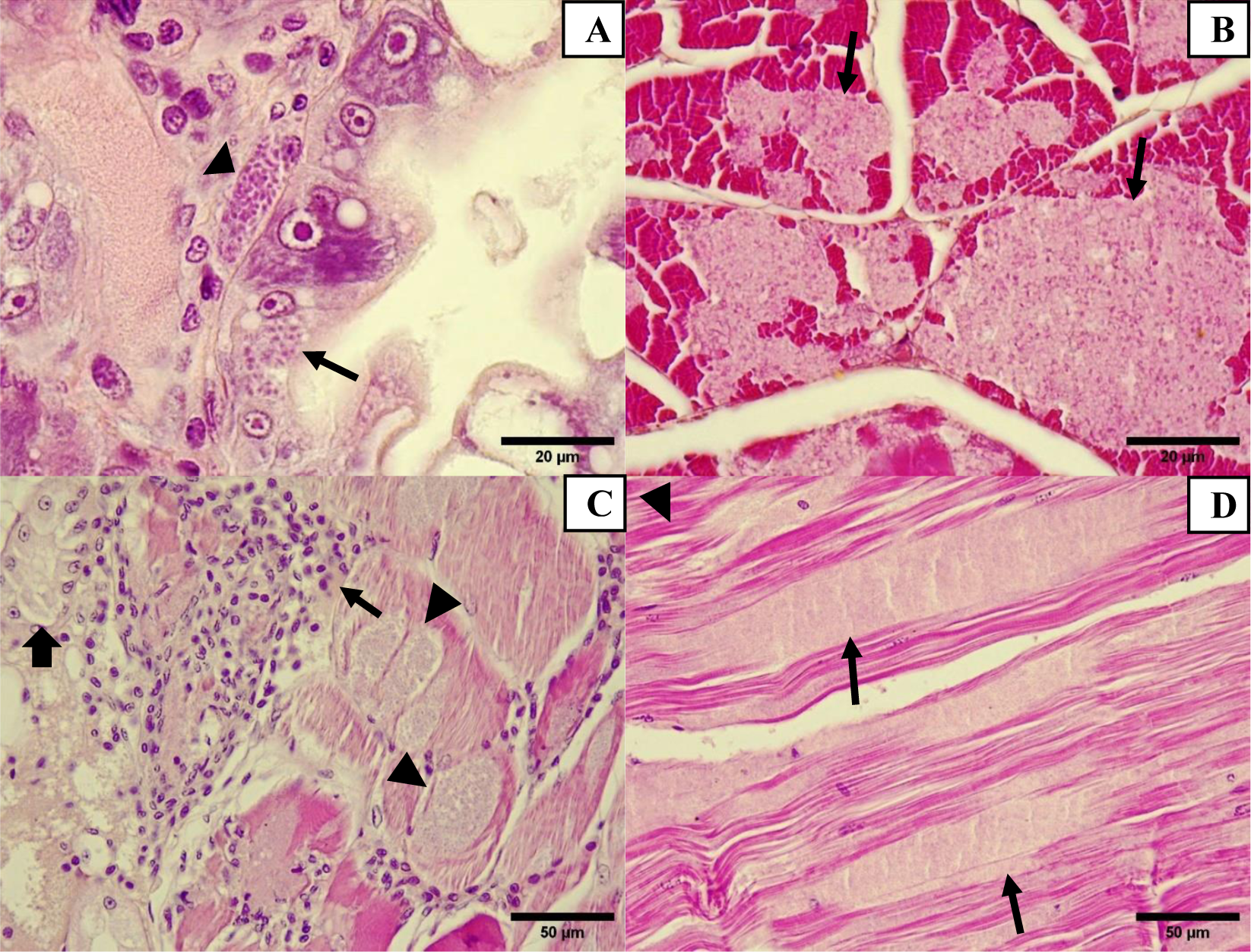
(A) Hepatopancreas of *P. vannamei* with microsporidia. The spores are in hepatopancreatic cells (arrow) and sinusoid space of hepatopancreas (arrowhead), 100X Magnification. (B) Cross-section muscle of *P. vannamei* showing spores-packed sarcomeres (arrows). (C) Anterior cephalothorax with and spore-packed sarcomeres, (arrowheads), adjacent to an antennal gland tubule (thick arrow). (D) Cephalothorax dorsal muscle with spore-packed (arrows) sarcomeres, H&E Stain.

**Pic 12:**
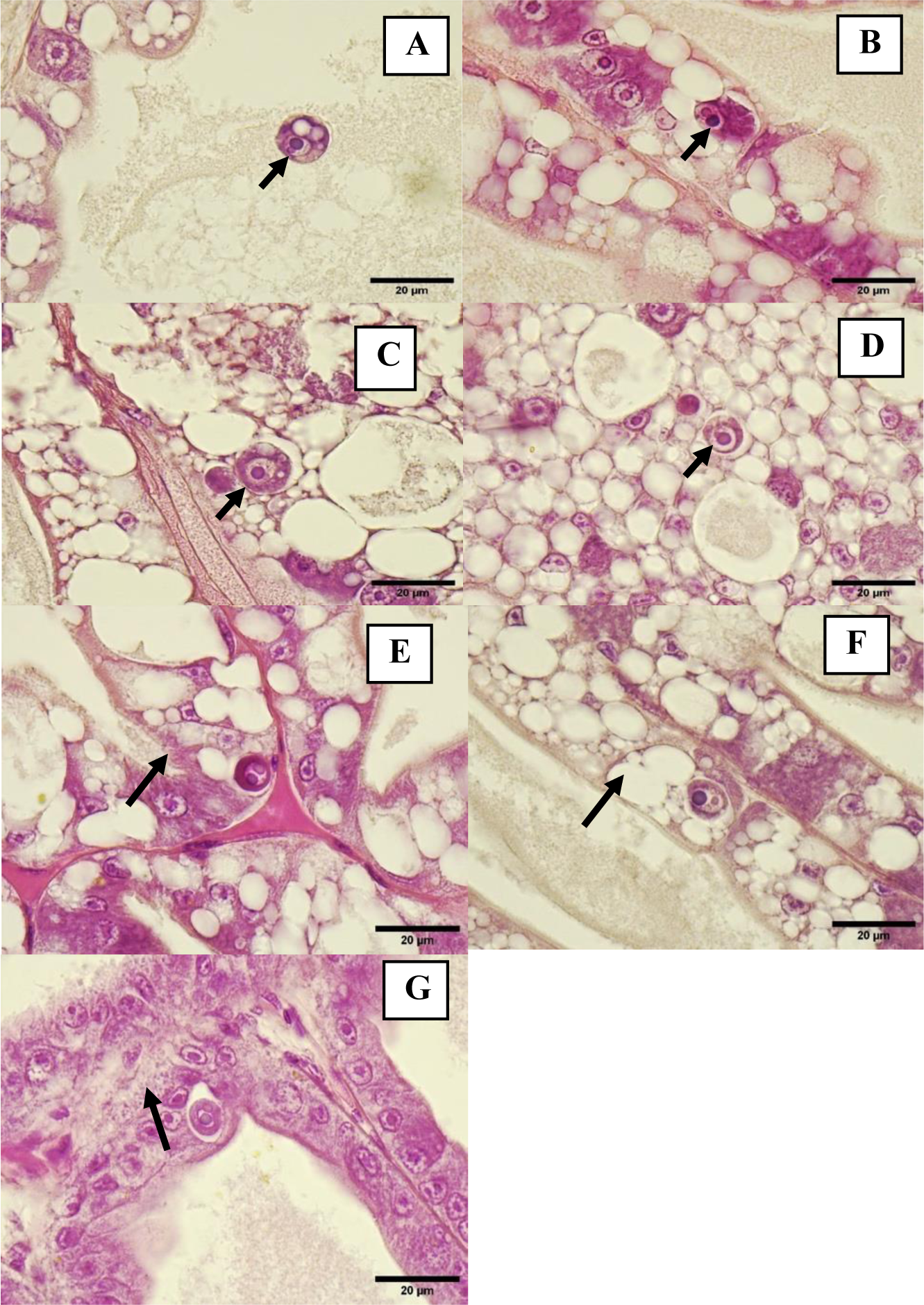
(A, B, C, D, E, F, G) Hepatopancreatic cells with viral inclusion body of WzSV8, inclusion body with double Lightner (arrow) and basophilic inclusion stage (arrowhead), H&E Stain.

**Pic 13:**
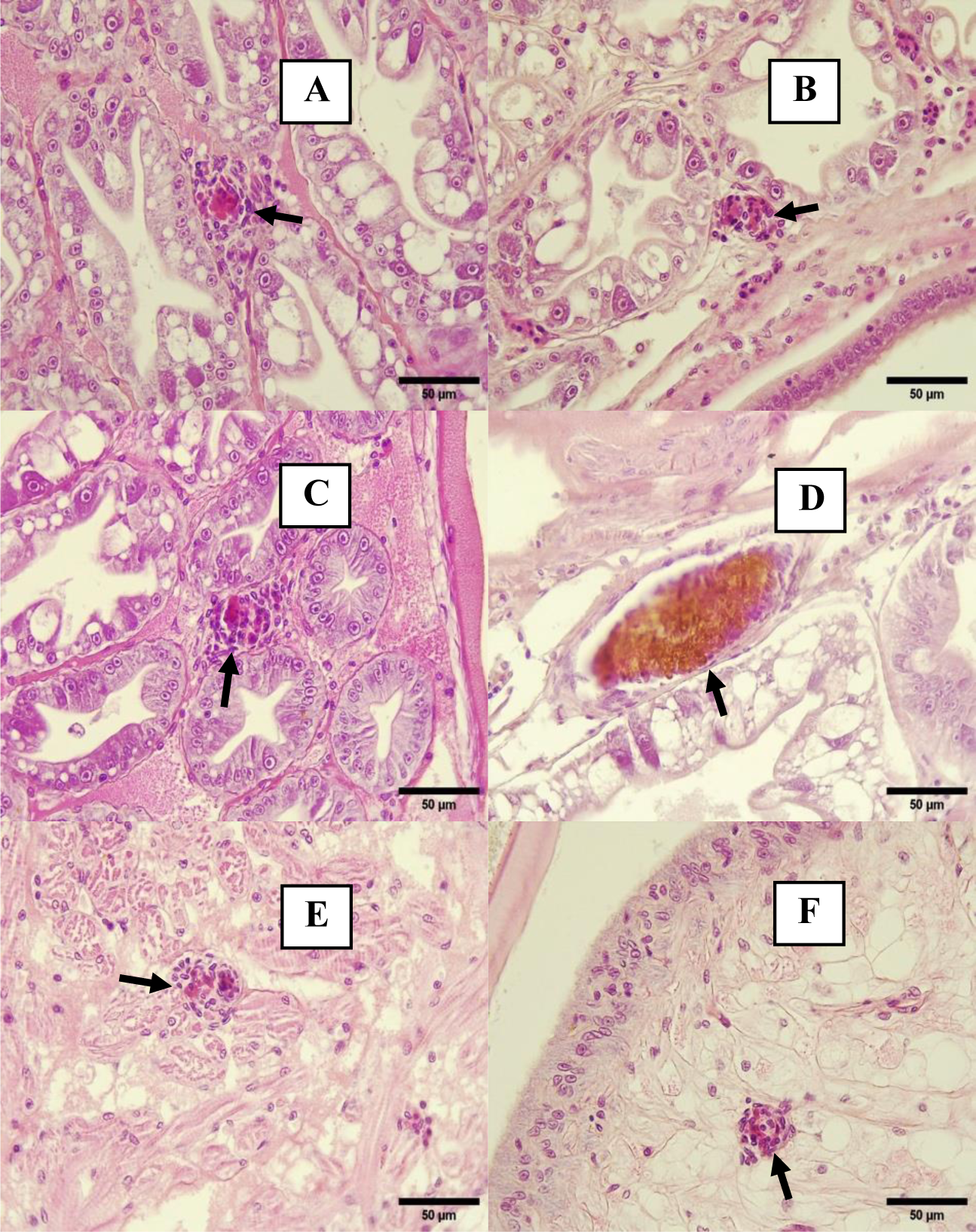
(A, B, C) Sinusoidal space of hepatopancreas with melanized nodule formation (arrow), 40X Magnification, H&E Stain. (D) Connective tissue next to hepatopancreas with melanized reaction (arrow), 40X Magnification, H&E Stain. (E) Cardiac muscle of heart with nodule formation (arrow), F) Connective tissue of stomach with nodule formation (arrow).

**Pic 14:**
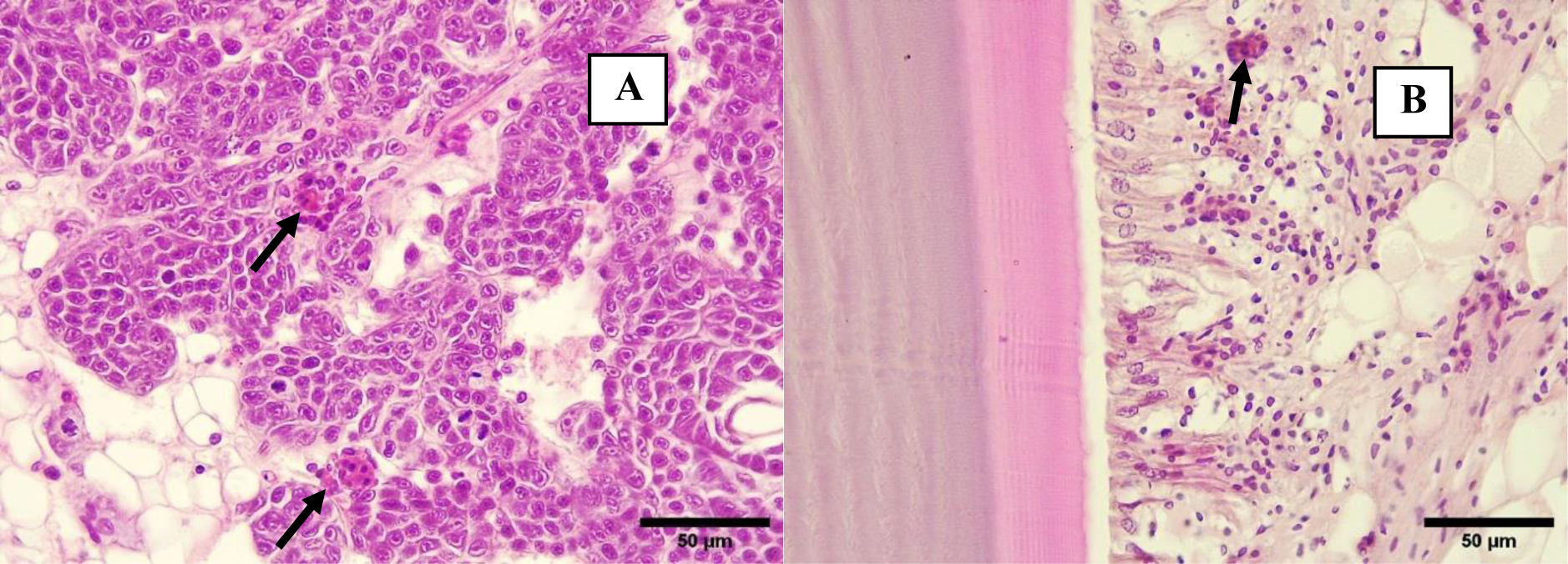
(A) Hematopoietic tissue with nodule formation (arrows). B) Epidermis and subdermis of cuticle with hemocytes infiltration and small nodule formation (arrow), H&E Stain.

**Pic 15:**
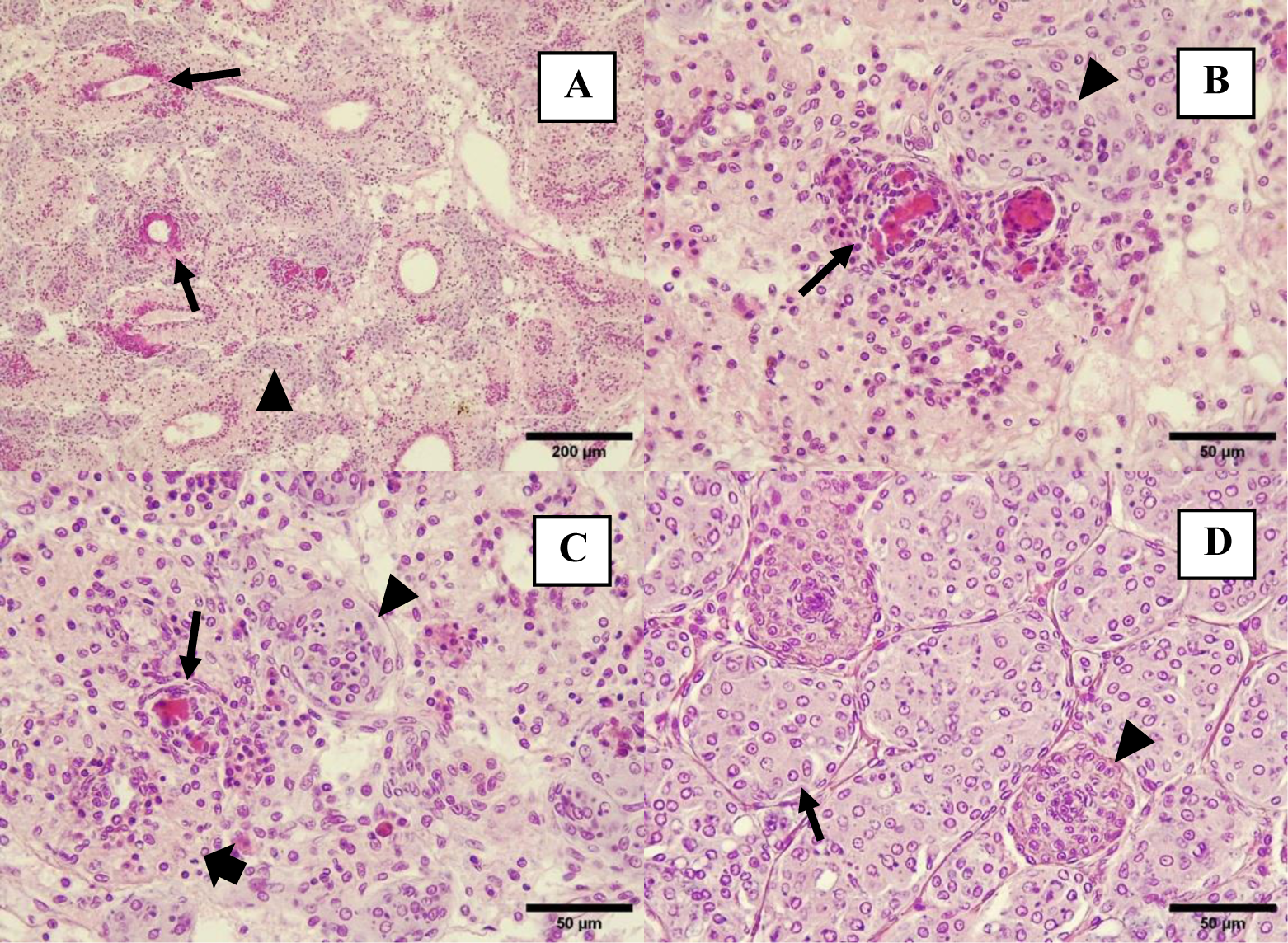
(A) Lymphoid organ with melanized reaction (arrow) and spheroid formation (arrowhead). (B, C) Lymphoid organ with melanized nodule formation (arrows), spheroid formation (arrowhead) and normal tubule (thick arrow), H&E Stain. (D) Lymphoid organ spheroid (arrow) and normal tubule (arrowhead), H&E Stain.

**Pic 16:**
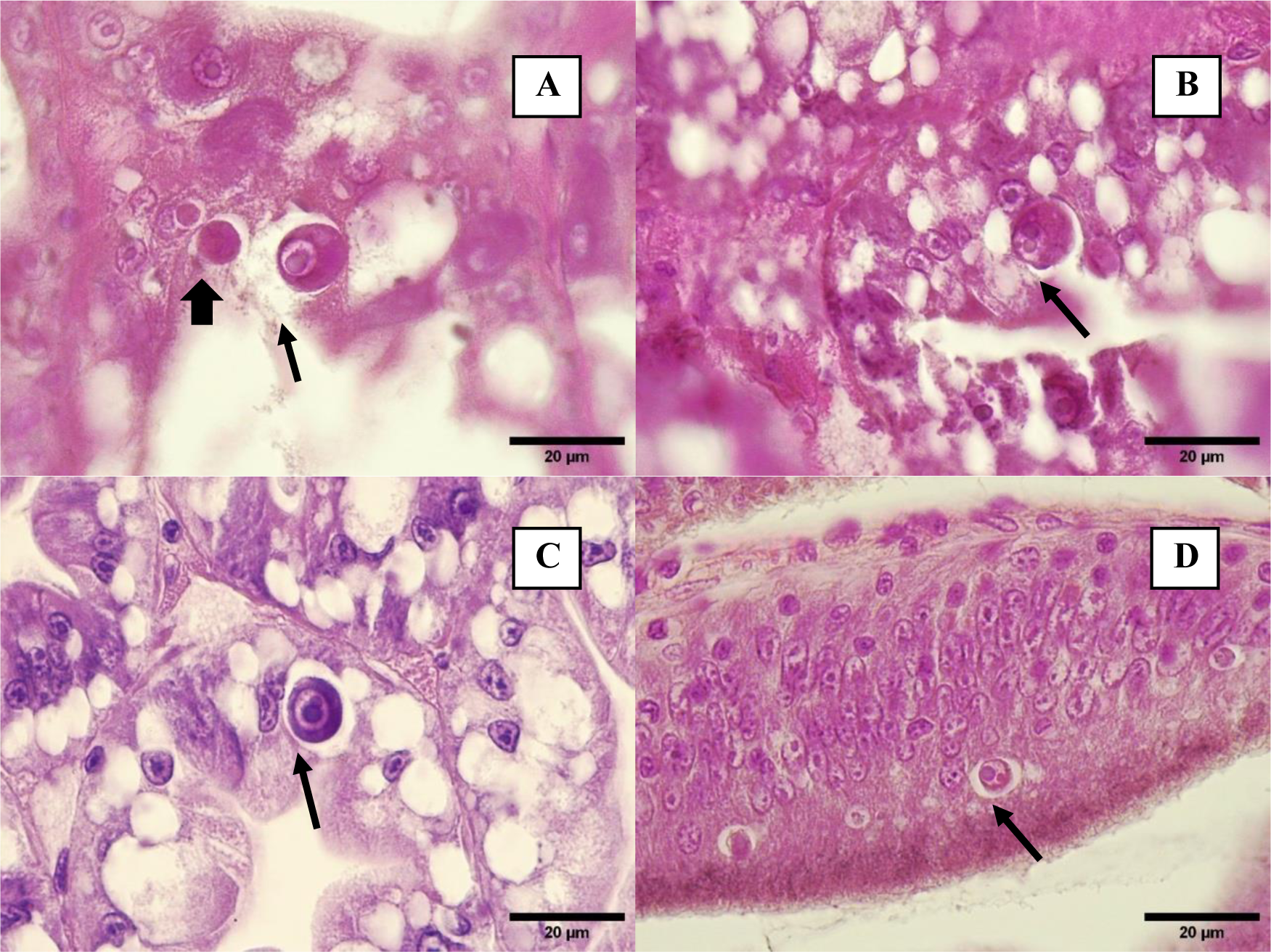
(A, B, C) Hepatopancreatic cells with viral inclusion body of WzSV8, inclusion body with double Lightner (arrow) and basophilic inclusion stage (arrowhead) (D) Epithelial cells of anterior midgut cecum with WzSV8 inclusion body. H&E Stain

**Pic 17:**
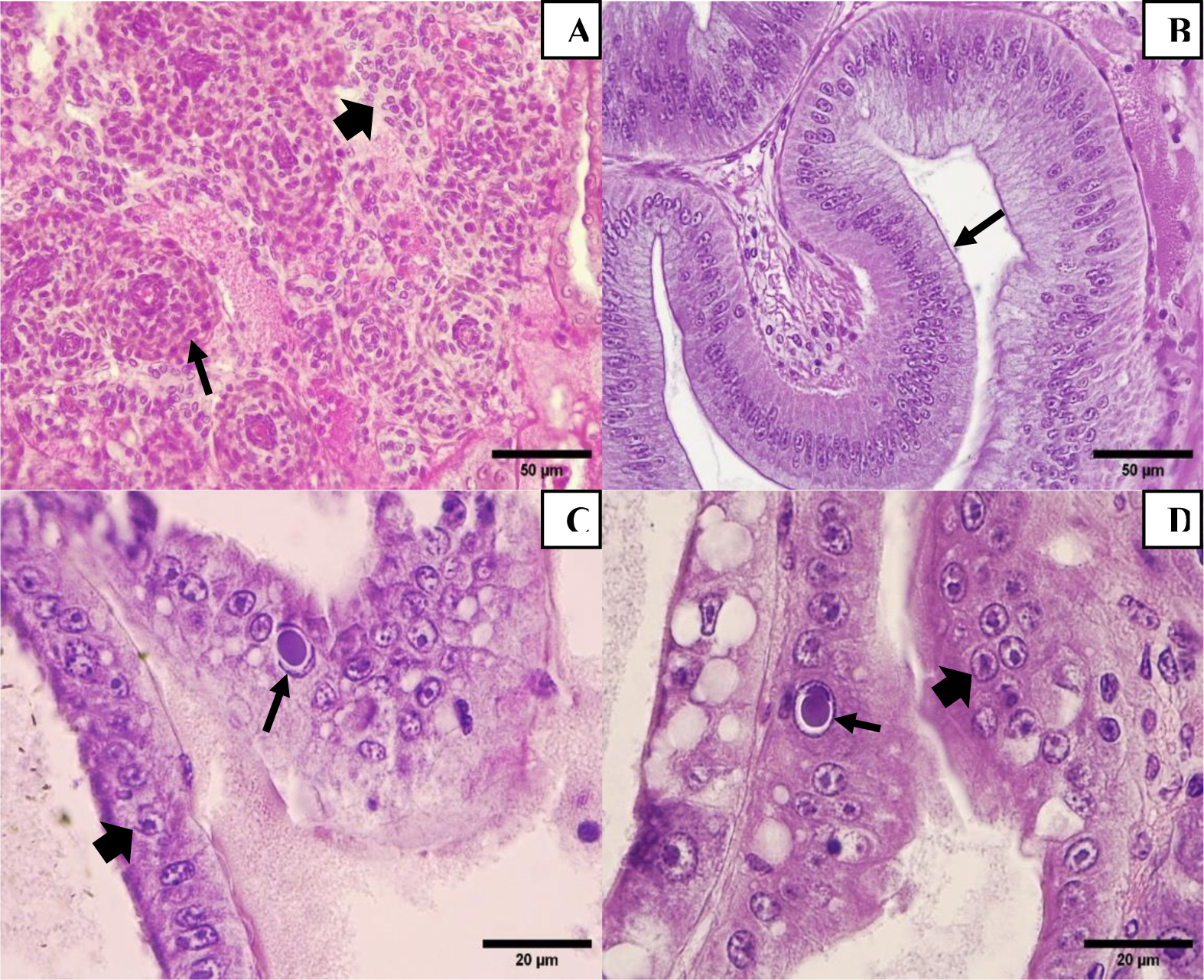
(A) Initial formation of LOS (arrow) and normal tubule of Lymphoid organ (Thick arrow), 40X. (B) Normal epithelium of midgut cecum. (C, D) Hepatopancreatic cells infected with DHPV (arrow), normal E-cells of hepatopancreas (thick arrow), H&E Stain

**Pic 18:**
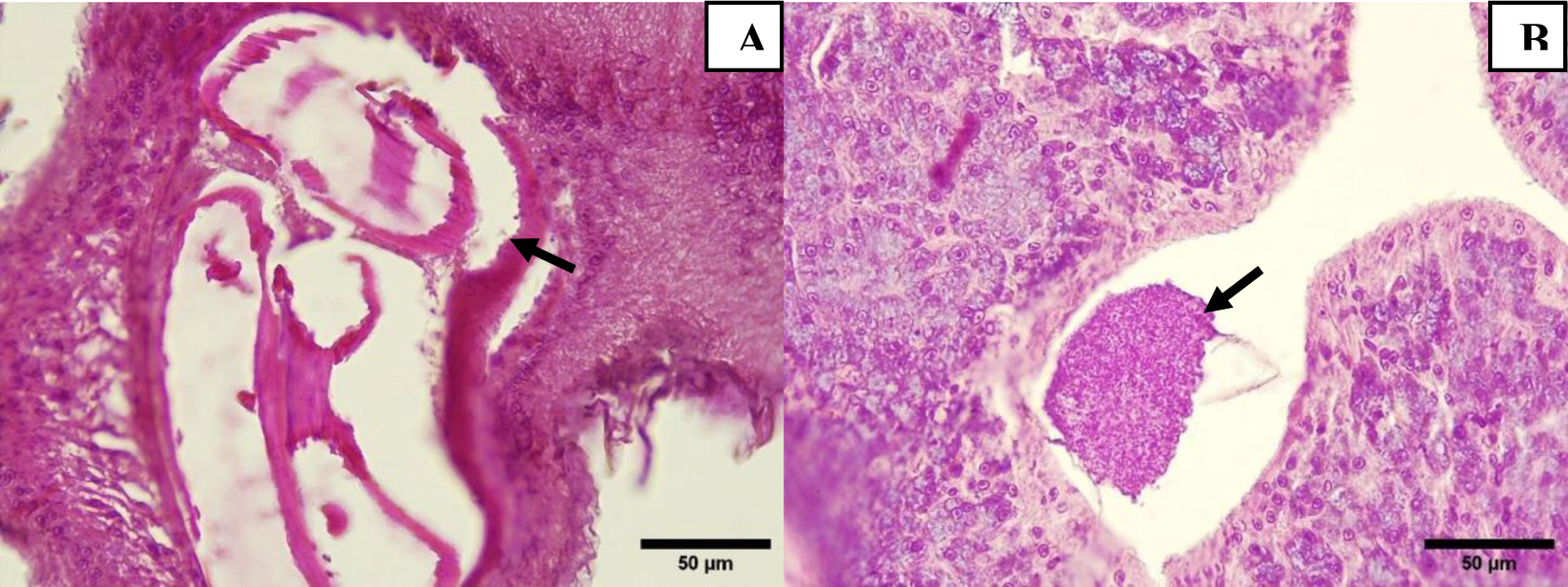
(A) Connective tissue of foregut with presence of a nematode cyst (arrow). (B) Lumen of Tegumental gland of hindgut caecum with presence of gregarine gametocyst (arrow), H&E Stain.

**Pic 19:**
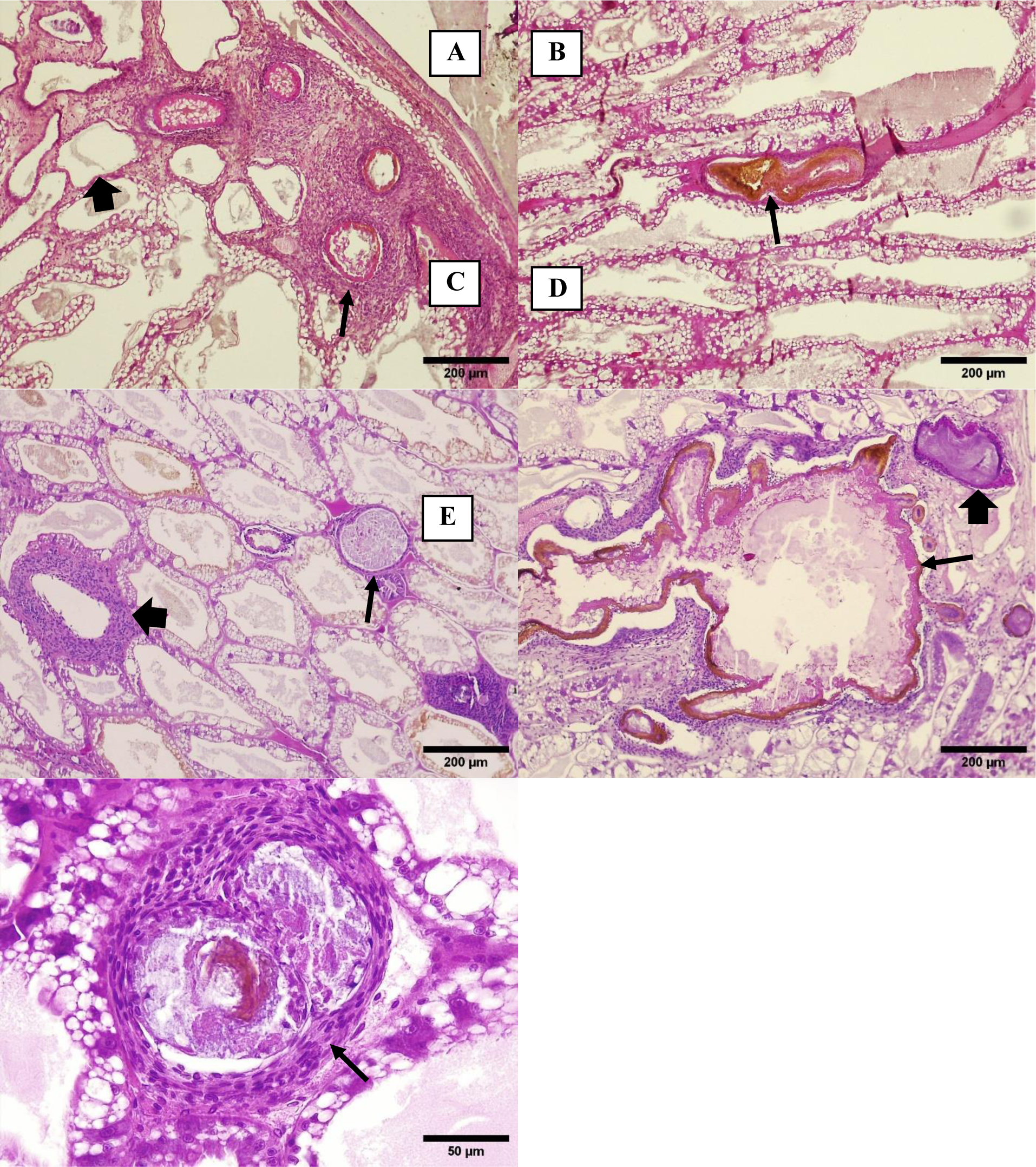
A) Hepatopancreas with area (arrow) of necrotic tubules and prominent host inflammatory response in the surrounding sinusoids (thick arrow): a tubule with atrophied epithelium. Note adjacent tubules appear to have normal R cell lipid vacuolation. (B) A single melanized tubule surrounded by tubules with heavily vacuolated epithelial cells. (C) Another view of necrotic tubules (thick and thin arrows) showing two different presentations of dysfunctional tubules, with adjacent tubules mostly well vacuolated. (D) Large area of consolidated, necrotic tissue and debris within a melanized capsule (arrow). An extension of the lesion (thick arrow). A necrotic, encapsulated tubule with tissue debris and microorganisms (arrow). Note “normal appearing vacuolation of adjacent tubule epithelium. H&E stain.

## Notes

### Competing Interest Statement

The authors have declared no competing interest.

### Summary of Updates

Some tables were corrected, the references corrected and updated. Missing references were incorporated.

